# The ALS/FTD-related C9orf72 hexanucleotide repeat expansion forms RNA condensates through multimolecular G-quadruplexes

**DOI:** 10.1101/2023.01.31.526399

**Authors:** Federica Raguseo, Anouk Huyghebaert, Jessica Li, Rubika Balendra, Marija Petrić Howe, Yiran Wang, Devkee M. Vadukul, Diana A. Tanase, Thomas E. Maher, Layla Malouf, Roger Rubio-Sánchez, Francesco A. Aprile, Yuval Elani, Rickie Patani, Lorenzo Di Michele, Marco Di Antonio

## Abstract

Amyotrophic lateral sclerosis (ALS) and frontotemporal dementia (FTD) are neurodegenerative diseases that exist on a clinico-pathogenetic spectrum, designated ALS/FTD. The most common genetic cause of ALS/FTD is the expansion of the intronic hexanucleotide repeat (GGGGCC)*_n_*in *C9orf72*. Here, we investigated the formation of nucleic-acid secondary structures in these expansion repeats, and their role in generating condensates characteristic of the diseases. We observed significant aggregation of the hexanucleotide sequence (GGGGCC)*_n_*, which we associated to the formation of multimolecular G-quadruplexes (mG4s), using a range of biophysical techniques. Exposing the condensates to G4-unfolding conditions led to prompt disassembly, highlighting the key role of mG4-formation in the condensation process. We further validated the biological relevance of our findings by demonstrating the ability of a G4-selective fluorescent probe to penetrate *C9orf72* mutant human motor neurons derived from ALS patients, which revealed clear fluorescent signal in putative condensates. Our findings strongly suggest that RNA G-rich repetitive sequences can form protein-free condensates sustained by multimolecular G-quadruplexes, highlighting their potential relevance as therapeutic targets for *C9orf72* mutation related ALS and FTD.

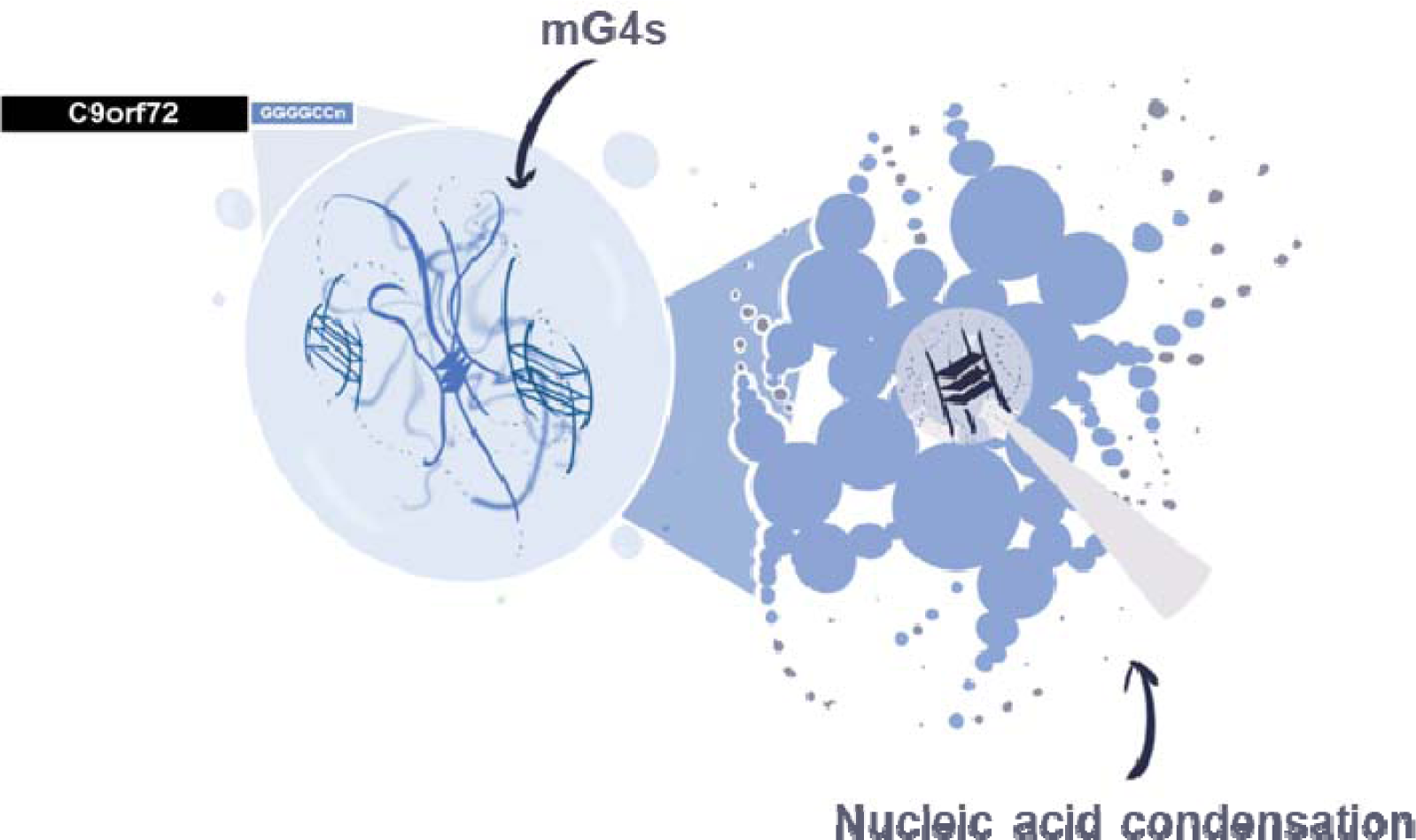

## Introduction

ALS is rapidly progressive, uniformly fatal and untreatable due largely to an incomplete understanding of disease mechanisms. The lifetime risk of ALS is 1:300-1:400, and over an aggressive disease course, patients become paralyzed, unable to eat, speak and breathe with an average survival of between 3-5 years^1^. Frontotemporal dementia (FTD) is the second most common cause of dementia in patients less than 65 years old and is increasingly recognized to share clinical, genetic and pathomechanistic features with ALS, termed ALS/FTD^2^. ALS and FTD, much like other neurodegenerative diseases, are characterised by the presence of pathological aggregates in neurons. Many existing studies have predominantly focused on the protein component as the leading aggregation trigger^3^, neglecting the role of nucleic acids as a key driver in the aggregation process^4^. Indeed, it is well established in ALS and FTD that RNA-binding proteins such as TDP-43 and FUS possess low-complexity domains which are prone to aggregation, due to low-affinity interactions^5,6,7^. In turn, recruitment of RNA in the aggregates has often been regarded as a secondary step of indeterminate significance. The most common hereditary cause of ALS/FTD is expansion of the intronic hexanucleotide repeat (GGGGCC)*_n_* in *C9orf72*, accounting for about 40% of familial ALS cases (where the patient had a family history of ALS) and 7% of the sporadic cases (no family history of ALS) in Europe^8^. This mutation has been previously proposed to lead to the formation of aggregates *via* various mechanisms^9,10^, including RNA transcribed from the expansion repeat playing a structural role. However, a clear mechanism to support this hypothesis has yet to be described^11,12^.

The hypothesis that RNA expansion repeats can aggregate in the absence of additional proteins has gained significant traction in recent years^13,14,15,16^. Although endogenous nucleic acid sequences do not typically lead to aggregation, they have the potential to form multimeric networks through both canonical and non-canonical base pairing interactions^17^. For example, r(CAG)*_n_*and r(GGGGCC)*_n_*, both natural sequences related to neurodegenerative diseases (respectively spinocerebellar ataxia^18^ and ALS/FTD), phase-separate *in vitro* at a critical number of repeats, suggesting that they can form multimolecular structures^19^.

r(GGGGCC)*_n_*, in particular, has previously been shown to arrange into hairpin and G-quadruplex (G4) structures, both of which have exhibited potential involvement in disease progression^20^. Indeed, the candidacy of these structures as potential therapeutic targets is demonstrated by using small-molecule probes that bind to and stabilise their secondary structures, leading to amelioration of disease phenotypes^20,21,22,23,24,25^. In particular, the use of ligands to bind G4s has been shown to ameliorate ALS phenotypes in neuronal cells^20,24,26,27^ and targeting of the hairpin with a small molecule inhibited repeat-associated non-ATG (RAN) translation and subsequent generation of toxic dipeptide repeats from the *C9orf72* gene mutation^21^. However, none of these studies have fully clarified the role of the nucleic acid structures in the regulation of pathological aggregation, which is a pre-requisite to devising optimised therapeutic agents. Furthermore, controversy exists about which secondary structure, G-quadruplex or hairpin, is most relevant to disease progression, and how. In this study, we attempted to address this controversy by investigating the role of an alternative structure that can be formed by the (GGGGCC)*_n_* repeat, which has not been studied in the ALS/FTD context: the multimolecular G-quadruplex structure (mG4). We hypothesized that multimeric G-quadruplexes could provide the three-dimensional linkages by the formation of G-G base pairing, which is required to initiate biomolecular condensates in *C9orf72* mutation-related ALS and FTD.

Indeed, non-canonical G-G base pairing underpinning G4-formation can readily occur under physiological conditions^28,29,30^ (**Figure 1A**) and G4-formation has already been associated with cancer, neurodegenerative disease (ALS and FTD) and other rare-genetic diseases (Fragile X syndrome and Cockayne Syndrome)^31,32^. The hexanucleotide intronic repeat (GGGGCC)*_n_* in the *C9orf72* gene has been shown to form G4s both in its RNA and DNA form^33,34,35,36^ and the RNA repeat r(GGGGCC) has been visualised within cellular ALS aggregates^22^. In addition, TDP-43 and FUS proteins, two RNA-binding proteins (RBPs) present in pathological aggregates in ALS (TDP-43 in >97% of cases^37^) and FTD (TDP-43 in approximately 40% of cases^38^ and FUS in approximately 10% of cases^39^), have both shown G4-binding abilities, reinforcing the idea that G4s might play a key role in the formation of condensates typical of ALS/FTD^40,41,42^. Furthermore, the hypothesis that mG4s are key to pathological aggregation would imply that targeting the hairpin conformation would slow disease progression by locking the RNA repeat in the hairpin conformation, preventing aggregate-formation. Similarly, treatment with G4-ligands could promote formation of unimolecular G4s, also preventing formation of mG4s and thus reducing the size of condensates upon treatment with G4-ligands, which would reconcile the seemingly discordant observations in the literature.

**Figure 1:**
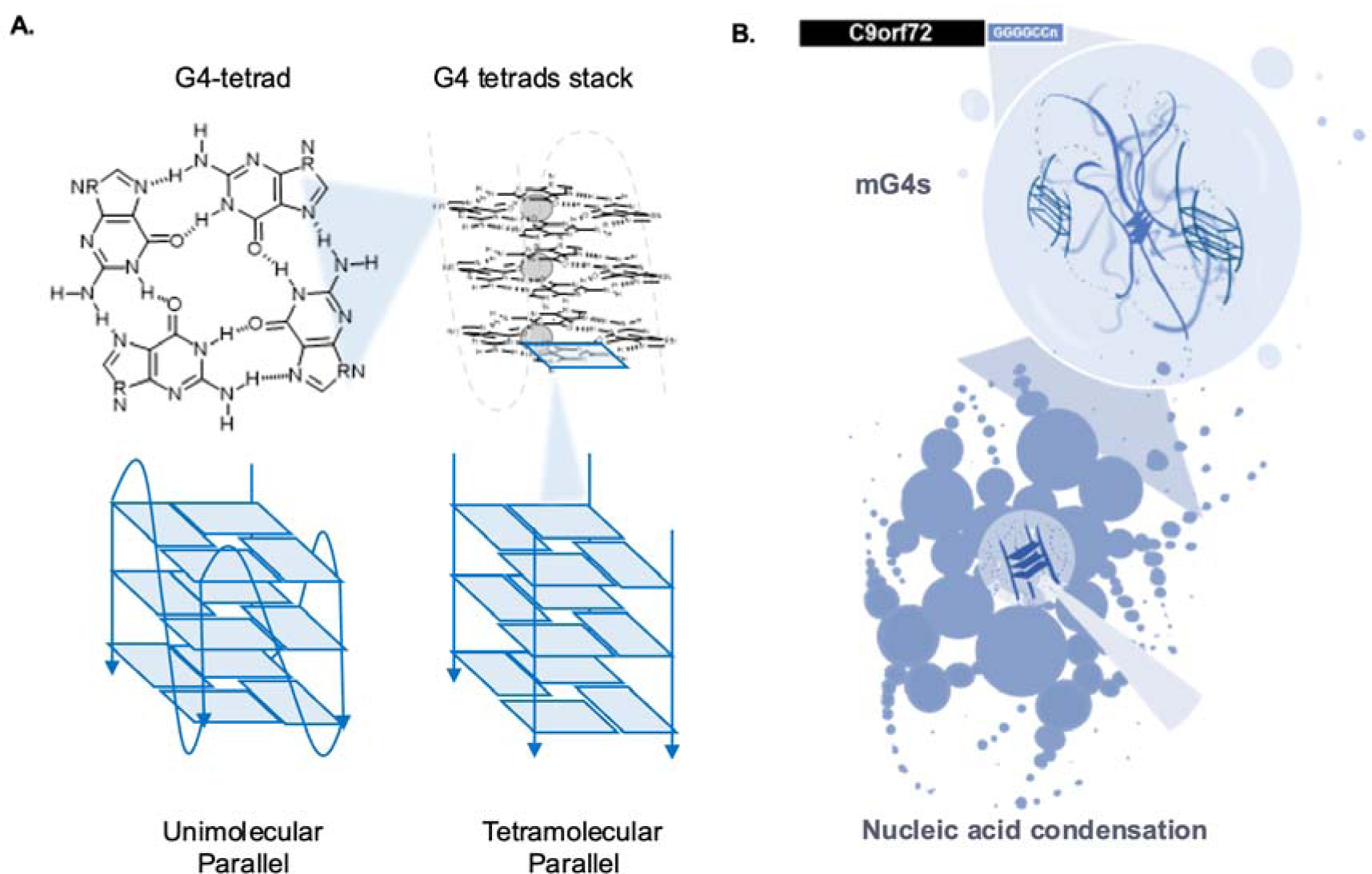
**A. Structural characteristics of G-quadruplexes -** The G4 scaffold is composed of a G-tetrad, a planar tetrameric assembly of guanine (G) bases linked together by Hoogsteen H-bonding. G4-tetrads stack to form a typical G4-structure, which is stabilised by monovalent cations sitting at the centre of the tetrad, the relative stability depending on the size of the cation (in order of increasing stability: Li^+^, Na^+^, K^+^), and more tetrads are held together by π-stacking interactions. **B. Proposed mechanism of aggregation in ALS and FTD pathological aggregates -** The (GGGGCC)*_n_* repeat sequence can aggregate in absence of proteins due to the formation of mG4s.

To investigate this, we extensively characterised the biophysical behaviour of (GGGGCC)*_n_* repeats to determine their ability to form mG4s and mG4-condensates under physiological conditions, observing substantial aggregation dependent on both repeat length *n* and oligonucleotide concentration. Next, we demonstrated that mG4s are structurally essential to condensation, as exposure to G4-unfolding conditions led to prompt condensate disassembly. Consistently, we observed that condensation is hindered by G4-ligand Pyridostatin (PDS), which has been previously suggested to favour binding to lower molecularity over multimolecular G4s^43^, and by rational design of mutants of the (GGGGCC)*_n_* repeat. These results suggest that the C9orf72 transcript itself aggregates through mG4s formation (**Figure 1B**). Furthermore, we have demonstrated that mG4 mediated condensation can be enhanced by the presence of the ALS/FTD related protein TDP-43, further indicating the potential for therapeutical relevance of our findings. Finally, we used the G4-selective fluorescent probe NMM to detect G4-structures in ALS-patient derived motor neurons. Cumulatively, our data provide mechanistic insight into pathogenic biomolecular condensation, a potential precursor to insoluble aggregate formation in ALS/FTD. Specifically, we demonstrate mG4 condensation in a protein-free environment and the interaction of mG4 with the most recognized ALS-related RBP TDP-43. Our findings may help to inform strategy for therapeutic targeting in currently these untreatable diseases.

## Results

To assess if multimolecular G4s (mG4s) can promote condensation of the hexanucleotide repeat sequence (GGGGCC)*_n_*, we started by investigating the ability of (GGGGCC)*_n_* to form mG4s and their stability to generate strong cross-links between nucleic acids strands.

### (GGGGCC)_n_ forms mG4s

We first investigated the ability of the (GGGGCC)*_n_* repeats to form multimolecular G4- structures. Although RNA is recognised as one of the toxic species in *C9orf72* mutation-related ALS and FTD pathological aggregates, we performed our initial biophysical studies with the DNA equivalents, as they are more easily sourced and handled^44^. To promote G4- formation, we annealed DNA (GGGGCC)*_n_* repeats of variable lengths (*n* =1 to 12) in a K**^+^**- containing buffer (500 mM KCl). To mimic intra-cellular crowding, 30% polyethylene glycol (PEG) was included in the buffer prior to slowly annealing the samples from 95 ℃ to 25 ℃ at a rate of -0.02 ℃/min (see methods for details).

Circular dichroism (CD) spectroscopy was initially performed to assess the formation of G4- structures. CD spectra of the (GGGGCC)*_n_* repeats annealed under mG4-forming conditions presented the characteristic negative peak at ∼240 nm and the positive peak at ∼263 nm, which are typical of a parallel G4-topology (**Figure 2A**)^45^. Although CD confirmed G4- formation, it cannot distinguish unimolecular from multimolecular topologies. Agarose gel electrophoresis (AGE) was thus employed to assess mG4s-formation (see methods for details, **Figure 2B**). Two main bands are observed, which are consistent with unimolecular and bimolecular species based on previous literature^32,35^ (**Figure 2B)**. It is noteworthy that the gel also indicated formation of bands at higher molecularity, which could not be visualised due to the low SYBR safe staining efficiency. To address this, we demonstrated formation of multimeric species that could be observed by AGE, using carboxyfluorescein (FAM)- labelled DNA (FAM) at lower concentrations to avoid smearing, as shown in **Figure S1**.

**Figure 2:**
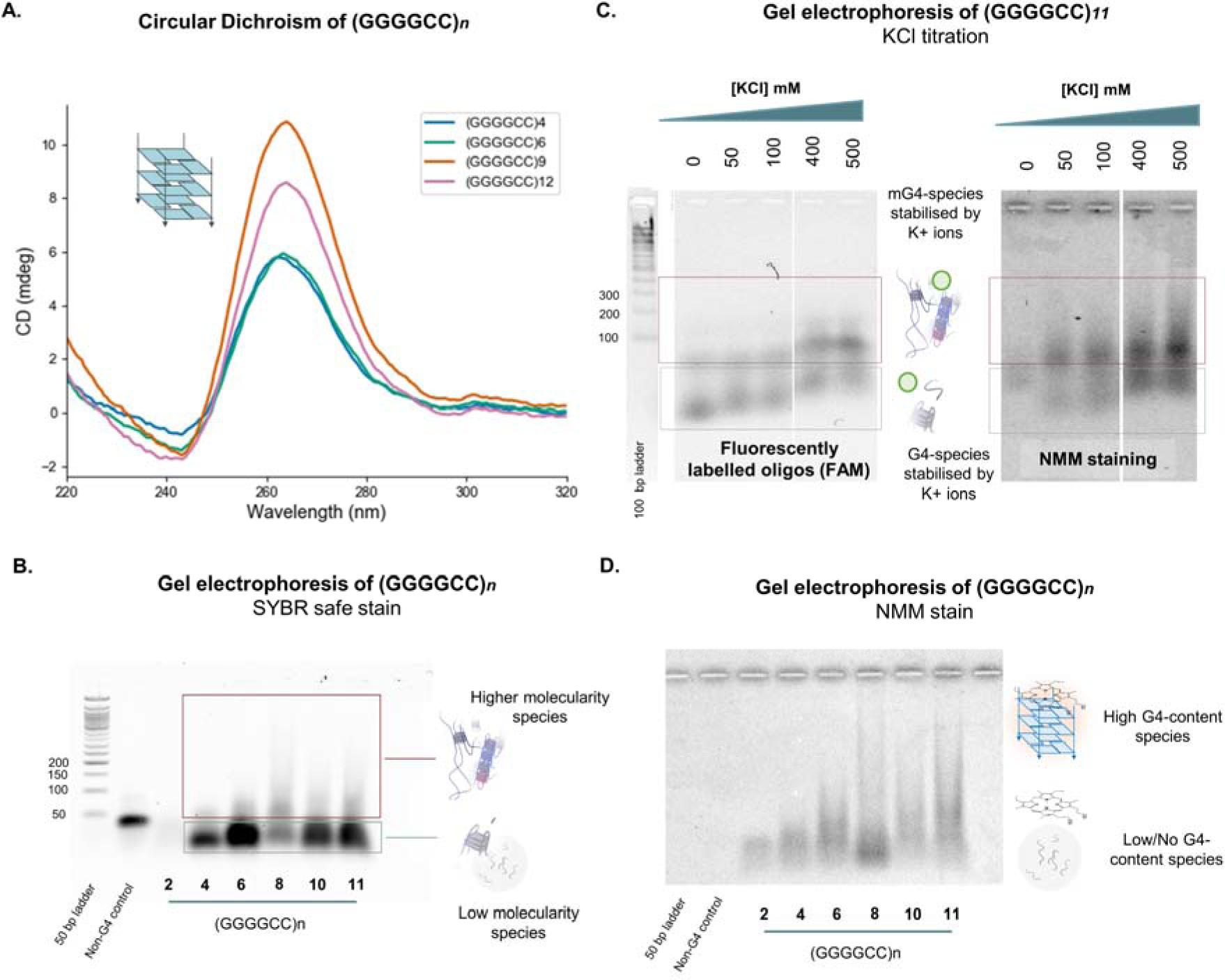
DNA (GGGGCC)*_n_* forms mG4s at different repeat lengths. **A.** (GGGGCC)*_n_* CD spectra - (GGGGCC)*_n_*(*n* = 4, 6, 9, 12) were annealed at 500 µM, 250 µM, 100 µM and 50 µM under mG4- forming conditions and further diluted to 5 µM for CD. All the samples present a positive peak at about 263 nm and a negative peak at 240 nm, both associated to a parallel G4 conformation. **B. (GGGGCC)*_n_* agarose gel electrophoresis – SYBR safe stain**. (GGGGCC)*_n_* (*n* = 2-11) were annealed at 250 µM in mG4-forming conditions and the gel was stained with SYBR safe. This gel includes the same samples used for the following NMM staining and has been added for reference to aid in the visualisation of the location of the bands. **C. (GGGGCC)_11_ agarose gel electrophoresis - (GGGGCC)_11_ was annealed in the presence of different KCl concentrations.** As KCl concentration increases, so does the molecularity of the species involved, indicating that the formed multimolecular structures have a strong KCl dependence. In the schematics, the green ball represents the K^+^ ion. On the right, the samples are stained with NMM, while on the left the sequences are FAM labelled. For reference a 100 base pairs ladder has also been added. **For visualisation, the ladder was stained with SYBR safe. D. (GGGGCC)*_n_* agarose gel electrophoresis – NMM stain.** (GGGGCC)*_n_* (*n* = 2-11) were annealed at 250 µM in mG4-forming conditions. The lanes containing any of the (GGGGCC)*_n_*were fluorescent upon staining with NMM, implying the presence of G4s in the higher molecularity species formed under these conditions. The absence of staining for both the ladder and the non-G4 ssDNA control confirm the specificity of the dye for G4-containing species.

The combination of CD and gel electrophoresis results suggested the ability of (GGGGCC)*_n_*sequences to fold into parallel G4s that can also adopt a multimolecular stoichiometry. To further demonstrate that the multimolecular species observed in the agarose gel could be ascribed to mG4s, we performed a KCl titration (0-500 mM) experiment, as multimolecular G4-formation is highly dependent on the concentration of K^+^. AGE experiments revealed that increasing K^+^ concentrations can quantitatively convert the fast-running band observed in **Figure 2B**, which we ascribed to the unimolecular G4, to a slowly running dimeric band, which is consistent with the formation of a bimolecular G4-structures (**Figure 2C**).

To further verify that the slow running bands observed by AGE contained mG4s, we also stained the samples with N-methyl mesoporphyrin IX (NMM), a G4-specific fluorescent dye^46,47,48^. NMM is expected to fluoresce upon binding to the bands containing G4s but not the non-G4 DNA control. Indeed, NMM selectively stains the slowly migrating portions of the (GGGGCC)*_n_*bands, indicating that the higher molecular weight bands contain G4s, as displayed in **Figure 2D**. To corroborate this observation, the AGE experiment was repeated using NMM staining, which confirmed the presence of G4s in the slow-moving band. The intensity of NMM fluorescence recorded in the slow-moving bands increases with K^+^, consistent with a greater abundance of (multimolecular) G4 species (**Figure 2C**).

### mG4s lead to DNA-condensation in absence of proteins

Having demonstrated the formation of multimolecular G4s in DNA samples of (GGGGCC)*_n_*, that increases in size with K^+^ concentration and length of the repeat, we proceeded to systematically screen for the emergence of macroscopic condensates formed by these repeats using confocal microscopy (see methods for mG4-annealing protocol conditions). **Figure 3A** shows representative images of the aggregates generated for different repeat lengths at a fixed concentration of 250 µM. Consistent with the observed clinical correlation between repeat length and pathological condensation in ALS and FTD^8^, we observed that *in vitro* condensation is more prominent for longer repeats. The condensates showed no fluorescence recovery after photobleaching (FRAP), indicating a solid or gel-like state (**Figure S2C**).

**Figure 3:**
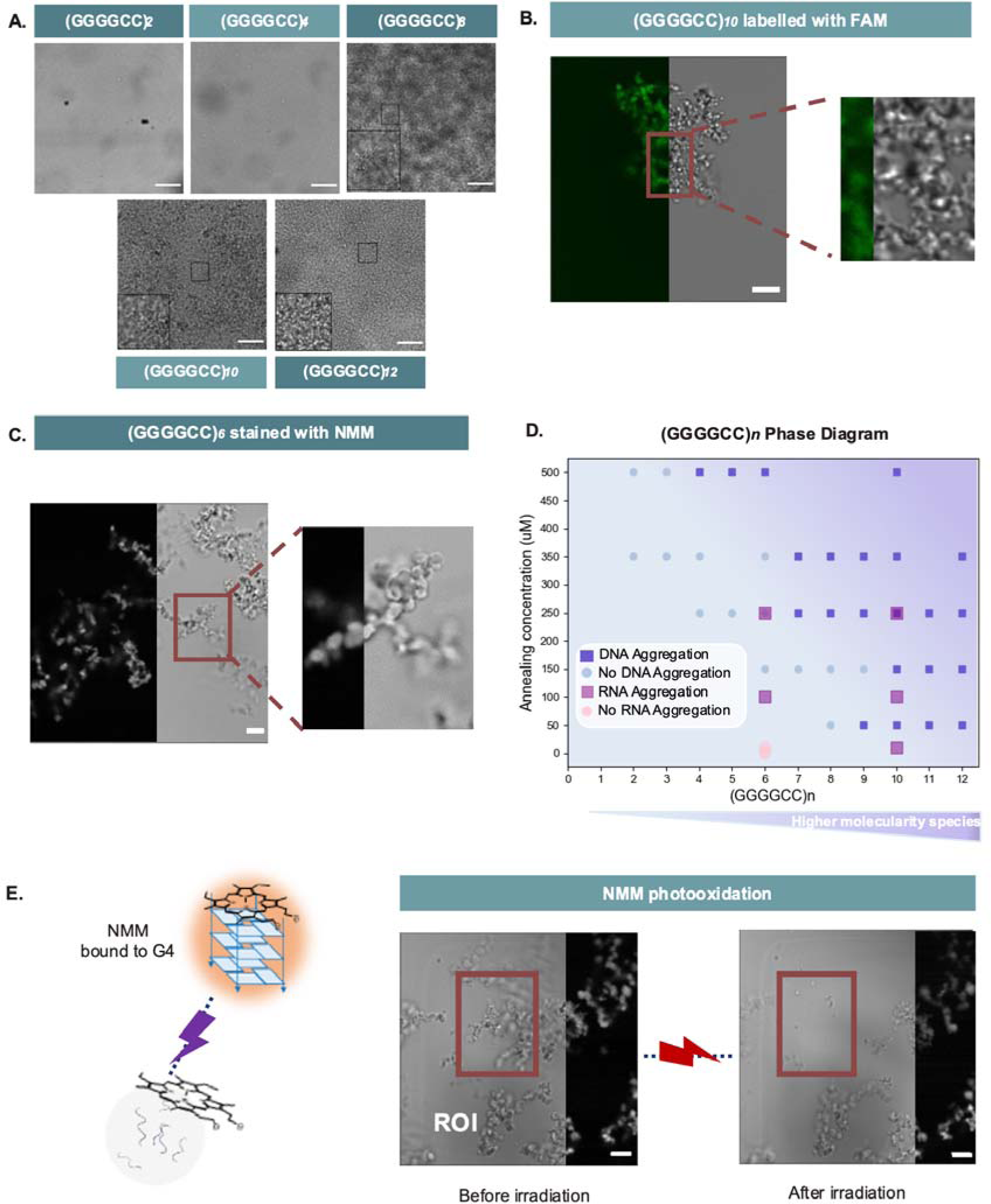
DNA (GGGGCC)_n_ aggregates in a G4-dependent fashion. **A. (GGGGCC)_n_ aggregates –** Brightfield imaging of (GGGGCC)*_n_* (*n* = 2-12) annealed under mG4-forming conditions at 250 µM. 100 µm scalebar. **B. FAM labelled (GGGGCC)_10_–** FAM-labelled (GGGGCC)_10_ was annealed in mG4- forming conditions. The image is split to show the brightfield channel (on the right) and the FAM (on the left). 10 µm scalebar. **C. (GGGGCC)_6_ NMM staining experiment** – (GGGGCC)_6_ was annealed under mG4-forming conditions and stained with NMM. The image is split to show the brightfield channel (on the right) and the NMM fluorescence channel (on the left). **D. (GGGGCC)_n_ phase diagram in KCl** – (GGGGCC)_n_ was annealed under mG4-forming conditions. At lower annealing concentrations and lower repeat lengths, the sequence does not present the ability to aggregate. The higher the repeat length and the higher the concentration, the more significant the aggregation. r(GGGGCC)_6,10_ were also annealed under mG4s forming conditions and presented aggregation at lower concentrations than their DNA equivalents. **E. (GGGGCC)_6_ NMM photooxidation experiment** – Upon laser excitation the NMM dye bound to the mG4s photo-oxidises the guanine in the quadruplex, leading to the disassembly of the structure. The region of interest highlighted in the figure (ROI) was irradiated for 1 minute and the second image was subsequently acquired. The aggregate in the ROI disassembled due to photooxidation. The image is split to show the brightfield channel (on the left) and the NMM (on the right). 10 µm scalebar.

The phase diagram in **Figure 3D** and **Figure S3** systematically maps the condensation state of the system as a function of repeat length and oligonucleotide concentration. Strands with repeat lengths of 2 and 3 showed no condensation even at the highest tested concentration of 500 µM. The shortest repeat-length producing condensates at 500 µM is *n* = 4, while constructs with *n*≥9 aggregated at oligonucleotide concentrations as low as 50 µM. The strong dependence of the phase-boundary location on repeat length confirms the positive correlation that an increased repeat length has on the formation of multimeric structures, consistent with AGE data in **Figures 2B** and **D**.

Microscopy results thus confirm that (GGGGCC)*_n_* are capable of forming condensates in the absence of proteins. Staining condensates formed by (GGGGCC)_6_ with NMM, expectedly produced a detectable fluorescence signal, indicating the presence of G4s in the condensates (**Figure 3C**). The selectivity of NMM-fluorescence for G4s aggregates over canonical dsDNA was assessed by staining a dsDNA condensate (**Figure S4**) under the same conditions. As expected, no fluorescence was detected in DNA condensates that lack G4s (**Figure S4B**).

Considering the data in **Figure 2**, we hypothesised that macroscopic aggregation was mediated by formation of mG4s causing the condensation of oligonucleotides. However, NMM staining does not *per se* imply a structural role of quadruplexes but only demonstrates their presence within the condensate. To assess whether G4s play a structural role within the condensates, we performed NMM photo-oxidation experiments (see methods for details). NMM binds to G4s by end-stacking and, upon irradiation with a high-power laser, it triggers guanine photo-oxidation, leading to the formation of 8-oxo-guanine that causes the disassembly of the G4-structure^49^. This phenomenon has been previously observed to not occur for dsDNA^49^, making it a highly G4-specific disassembly route, which we have previously leveraged to achieve light-controlled disassembly of mG4s in engineered DNA nanostructures^50^. As shown in **Figure 3E**, we irradiated a portion of the (GGGGCC)*_6_*NMM- stained condensates to trigger photo-oxidation, which led to immediate disassembly of the condensates. Given that no other relevant multimolecular interaction is known to be affected by NMM photo-oxidation, these results suggest that mG4s are the prevalent structure holding the condensate together, and that in the absence of guanine-guanine hydrogen bonding, necessary for G4-formation, no aggregation is observed. Moreover, we tested the selectivity of the assay on NMM-stained dsDNA-condensates, which did not show any significant change upon irradiation (**Figure S4C**). Notably, although the samples analysed were annealed in 500 mM KCl to probe mG4 formation in higher yields, condensates were also observed at physiological KCl amounts for higher DNA concentrations (**Figure S5**).

Additionally, to ensure that the crowding agent was not triggering DNA condensation in its own right, (GGGGCC)_11_ annealing was also performed in absence of PEG with a slower cooling rate to probe mG4-formation. As shown in **Figure S6**, condensates were still able to form, confirming that the crowding agent was not influencing DNA condensation.

### (GGGGCC)_n_ mutants designed to prevent mG4-formation reduce aggregation

To further confirm that mG4s were responsible for condensation observed for (GGGGCC)_n_ repeats, we investigated the condensation properties of a series of mutated sequences (**Figure 4A**).

**Figure 4:**
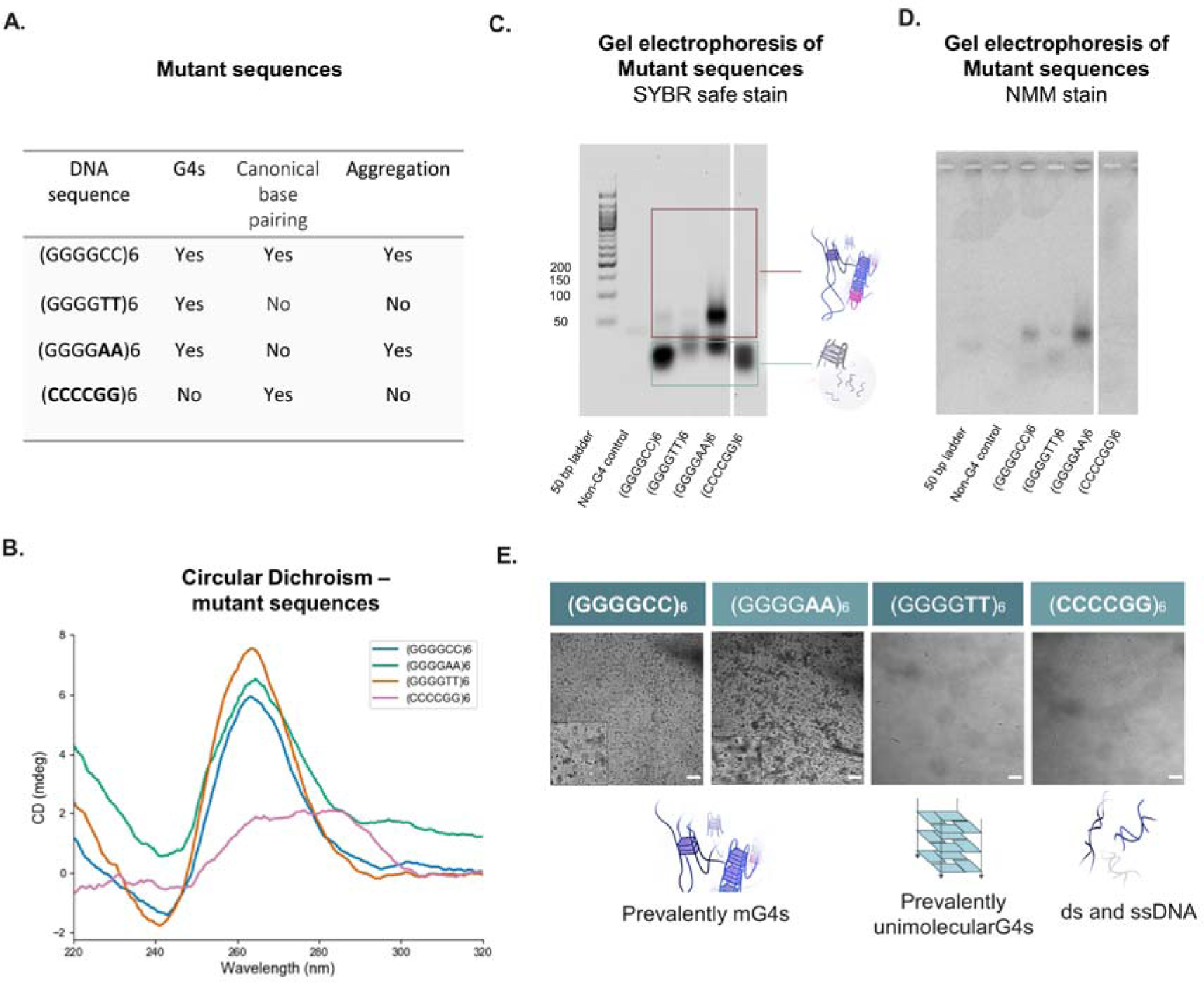
Role of base pairing and monomolecular G4s in the aggregation mechanism. **A. Table of mutants. B. Circular Dichroism spectra of mutants** – Controls of *n* = 6 were annealed in mG4-forming conditions. Only the predicted G4-forming sequences present the characteristic peaks at >260 nm and 240 nm. **C. Mutants agarose gel electrophoresis – SYBR safe stain** – Controls of *n* = 6 were annealed in mG4-forming conditions. D. (GGGGCC)*_n_*agarose gel electrophoresis – NMM stain – Controls of *n* = 6 were annealed in mG4-forming conditions. All G4-forming sequences ((GGGGAA)_6_, (GGGGTT)_6_, (GGGGCC)_6_) presented fluorescence upon staining, confirming the presence of the G4. Ladder, non-G4 ssDNA control, and CCCCGG_6_ do not appear, confirming the specificity of the dye for G4-containing species. **E. Aggregation of the mutant sequences** – Controls of *n* = 6 were annealed under mG4-forming conditions. Aggregates were only clearly distinguishable for (GGGGAA)_6_ and (GGGGCC)_6_ – the only sequences forming mG4s. 100 µm scalebar.

In addition to G4-formation it is possible that GC base pairing may play a role in stabilising the observed aggregates. To test this hypothesis, we studied the (CCCCGG)_6_ repeat, which retains the same number of potential canonical base pairing interactions as (GGGGCC)_n_ but is unable to form G4s. CD was initially performed to confirm the absence of G4-formation, which was further validated *via* NMM staining of these sequences loaded on an agarose gel (**Figures 4B**, **C** and **D**). Replacing Cs with As in the (GGGGAA)_6_ repeat preserves G4- forming ability but prevents the formation of canonical base-pairing. CD of this repeat revealed a typical parallel-G4 trace and NMM staining of the agarose gel confirmed G4 formation (**Figures 4B** and **C**). Similarly, (GGGGTT)_6_ maintains the ability to form G4s preventing base-pairing, but due to G-T stacking interactions it is expected to favour mono- molecular G4-structures over multimolecular ones^51^, allowing us to disentangle the effect of unimolecular vs multimolecular G4s in the aggregation process. CD and agarose gel-NMM staining of the T-to-C mutant indeed revealed formation of G4s (**Figures 4C** and **D**).

Having gauged the G4-forming tendencies of the mutants, we proceeded to assess their condensation properties. No significant condensation was observed for (CCCCGG)_6_, confirming that canonical base pairing interactions and hairpin formation are not sufficient to drive condensation of the oligonucleotides. Similarly, no significant condensation was observed for (GGGGTT)_6_, which predominantly forms unimolecular G4s as opposed to mG4s (see AGE in **Figure 4C**). This evidence further supports our hypothesis that mG4s are the essential cross-linking elements in the condensates, while formation of G4s *per se* is insufficient to prompt condensation. Consistently, (GGGGAA)_6_ and (GGGGCC)_6_ were the only two sequences to reveal significant mG4-formation in the AGE experiments, and to show formation of condensates under confocal microscopy (**Figure 4E**).

### PDS prevents (GGGGCC)_n_ aggregation

G4 ligands have been shown to ameliorate the presence of ALS aggregates in cells^26,52^. The ligands have been speculated to act by disrupting the toxic downstream functions of C9orf72 - thought to underpin aggregation^52^. Having demonstrated that mG4s alone are able to prompt aggregation in protein-free samples, we asked whether G4 ligands could destabilise the condensates by promoting the formation of unimolecular as opposed to multimolecular G-quadruplexes. For this purpose, we employed pyridostatin (PDS), a well-characterised G4-ligand^43^, known to form more stable complexes with lower molecularity G4s over multimolecular G4s^53^ and compatible with our experimental conditions.

NMM-stained AGE of (GGGGCC)_10_ annealed in the presence of 10 µM PDS revealed a substantial reduction in high molecularity species compared to the PDS-free sample (**Figure S7**). The ability of PDS to suppress large multimers is reflected by a substantial reduction in the number and size of condensates visible by confocal microscopy (**Figure 5**; see methods for details). It is worth noting that the small percentage of DMSO introduced to solubilise the ligand does not directly impact aggregation.

**Figure 5:**
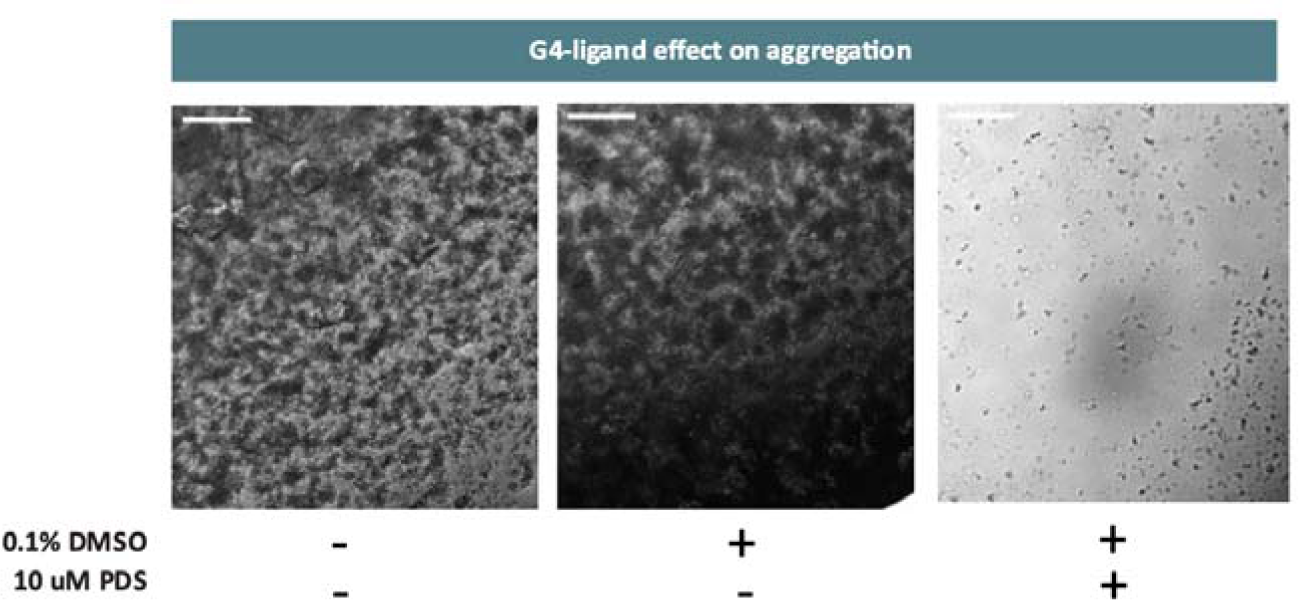
(GGGGCC)_10_ aggregation in presence of 10 µM PDS. – (GGGGCC)_10_ was annealed at 150 µM under mG4-forming conditions. The sample containing the G4-ligand presents aggregates to a far lesser extent with respect to the non-PDS containing controls. 100 µm scalebar.

Beyond further confirming the critical importance of G4 molecularity in stabilising (GGGGCC)_n_ aggregates, these data outline a potential new mechanism for the reported ability of G4 ligands to ameliorate reported ALS phenotype in cells^26^.

### RNA repeats form biomolecular condensates at lower concentrations than their DNA counterparts

RNA is considered a candidate toxic species in *C9orf72* mutation-related ALS and FTD pathological aggregates. As DNA equivalents were tested in the preceding experiments, we sought to address whether RNA repeats conformed to similar behaviour and whether any notable differences existed. Indeed, RNA G4s (rG4s) are known to arrange preferentially into a parallel conformation and to be more thermodynamically stable than the equivalent DNA G4s^54,55^. For these reasons, we hypothesised that aggregation of r(GGGGCC)*_n_*occurs at lower concentrations in RNA compared to with DNA but that, overall, the aggregates would possess similar physical and chemical properties.

Therefore, we sought to confirm the ability of RNA to form condensates in a similar manner to its DNA equivalent by subjecting r(GGGGCC)_6/10_ to the same condensation conditions. Aggregation was observed for concentrations as low as 100 µM for repeat length 6, and 10 µM for *n* =10 (**Figure S8**), forming amorphous aggregates similar in morphology to their DNA equivalents under all conditions. For comparison, DNA (GGGGCC)_6_ was seen to aggregate only above 350 µM in concentration, reflecting the known ability of RNA to form more stable G4s compared to DNA. To further validate mG4-formation in the RNA aggregates, samples were stained with NMM, confirming the presence of G4s (**Figure 6A**).

**Figure 6:**
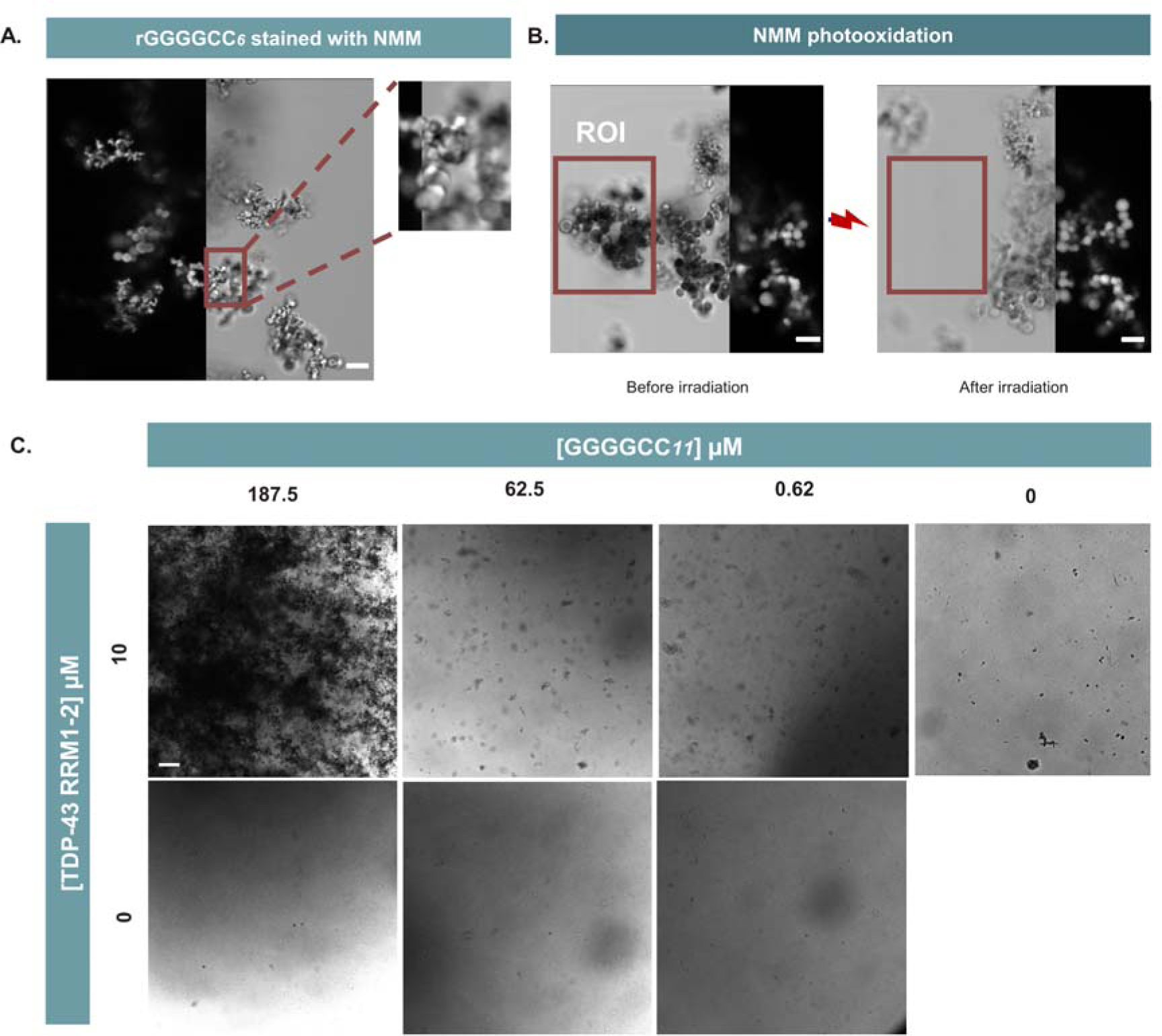
A. RNA (GGGGCC)*_n_*aggregates with a G4-dependent trend, r(GGGGCC)*_6_* NMM staining experiment – r(GGGGCC)_6_ was annealed in mG4-forming conditions. It was then subjected to NMM staining prior to acquisition. The image is split to show the brightfield channel (on the right) and the NMM fluorescence channel (on the left). **B. RNA (GGGGCC)*_n_* aggregates with a G4- dependent trend,** r**(GGGGCC)*_6_* NMM photo-oxidation experiment –** Upon laser excitation the NMM dye bound to the mG4s photo-oxidises the guanine in the quadruplex, leading to the disassembly of the structure. The region of interest highlighted (ROI) in the figure was irradiated for 1 minute and the second image was subsequently acquired. The aggregate in the ROI disassembled due to photo- oxidation. 10 µm scalebar. **C. Brightfield imaging of TDP-43 RRM1–2 and (GGGGCC)_11_ NMM- stained aggregates -** (GGGGCC)_11_ was annealed under mG4-forming conditions (1-300 µM) and then incubated with 10 µM TDP-43 RRM1–2. Upon incubation with the protein, the mG4 aggregates trigger protein, indicating a synergist aggregation effect between the (GGGGCC)*n* repeats and TDP-43. 100 µm scalebar.

Furthermore, the RNA aggregates stained with NMM were subjected to irradiation at 405 nm to induce guanine photo-oxidation, which caused immediate disassembly of the irradiated aggregates (**Figure 6B**), as previously observed for DNA. These results confirmed that (GGGGCC)*_n_* aggregation is an mG4-dependant phenomenon and occurs in a similar manner for RNA and DNA repeats, but crucially requires lower concentrations in the case of RNA.

### (GGGGCC)_n_ enhances aggregation of TDP-43

Transactive response DNA binding protein of 43 kDa (TDP-43) is a nucleic acid-binding protein that plays a crucial role in RNA processing and regulation of gene expression and has been extensively studied in the context of neurodegenerative diseases, particularly in ALS/FTD where it forms pathological aggregates in affected neurons^56^. Interestingly, this protein has previously shown G4-binding abilities, raising the possibility that its interaction with G4s enhances or even triggers pathological aggregation^40^.

To test this hypothesis, we purified a TDP-43 protein variant which only comprises the RNA binding region, *i.e.*, RRM1–2 (K102–Q269)^57^ (**Figure S9**). We selected this variant because it binds to nucleic acids similarly to full length TDP-43. Furthermore, it is more soluble than full length TDP-43 *in vitro* after purification, yet it can form aggregates under specific conditions that resemble physiology^57^. The aggregation propensity of this variant was sufficient to accurately permit the assessment of protein and mG4 condensation over a range of concentrations. First, we assessed whether the protein could aggregate with (GGGGCC)_n_. We annealed FAM-labelled (GGGGCC)_11_ under mG4-forming conditions (100 µM) and incubated it for 3 days with 10 µM of TDP-43 RRM1–2 labelled with ALEXA 633. The resulting aggregates were active in both fluorescence channels (**Figure S10A**), suggesting that TDP-43 RRM1–2 was co-aggregating with the (GGGGCC)_n_ repeat. We then investigated how the presence of TDP-43 RRM1–2 impacts (GGGGCC)_11_ aggregation. To achieve this, we screened the aggregation properties for a series of (GGGGCC)_11_ concentrations (0,1,100,300 µM) in the presence or in the absence of a fixed concentration of TDP-43 RRM1–2 (10 µM).

As shown in **Figure 6C**, we detected more condensation events when the (GGGGCC)_11_ repeat was exposed to TDP-43 RRM1–2, suggesting that the protein domain can enhance the aggregation of the oligonucleotide. Importantly, TDP-43 RRM1–2 alone does not form aggregates that are visible at the limit of this microscope’s resolution, suggesting that the emergence of larger aggregates requires both the protein and nucleic acid components. This was subsequently confirmed via transmission electron microscopy studies where smaller TDP-43 RRM1-2 aggregates can be observed with a higher resolution (**Figure S11**), consistent with previous literature^58^. To confirm that mG4s were still present in the final protein/nucleic acids condensates, the samples were stained with the G4-specific fluorescent probe NMM. As shown by NMM staining condensates formed in the presence of TDP-43 RRM1-2 still had a significant G4-content (**Figure S8B**). Given the pathological relevance of TDP-43 aggregation in ALS and FTD, we anticipate that the observed synergy between RRM1–2 and (GGGGCC)_n_ repeats could bear significance for the development of future therapeutic strategies.

### ALS/FTD RNA aggregates can be visualised by NMM in C9orf72 mutant patient derived neurons

Encouraged by the enhanced TDP-43 aggregation observed in the presence of the (GGGGCC)_n_ aggregates, we sought to further validate the physiological relevance of our findings by using ALS/FTD patient-derived spinal cord motor neurons generated from induced pluripotent stem cells (iPSCs) carrying the *C9orf72* mutation. Here we used a fully established human stem cell model^59^, which has previously been reported to reveal RNA *foci* that can be visualised by microscopy^60,61^.

We sought to address whether RNA aggregates in the ALS/FTD phenotype contain G4- structures within them. To achieve this, we stained fixed human C9orf72 mutant neurons with NMM (10 µM) to visualise G4s within the aggregates and the nuclear dye DRAQ5 to identify the nuclei of the cells. Indeed, we observed positive NMM staining in putative aggregates, in this previously characterized human stem cell model^60^ **(Figure 7A)**. The occurrence of G4s in neurons can be also captured by flow cytometry (**Figure S12**), which showed a notably enhanced fluorescent intensity upon NMM (10 µM) treatment **(xFigure 8B)**.

**Figure 7:**
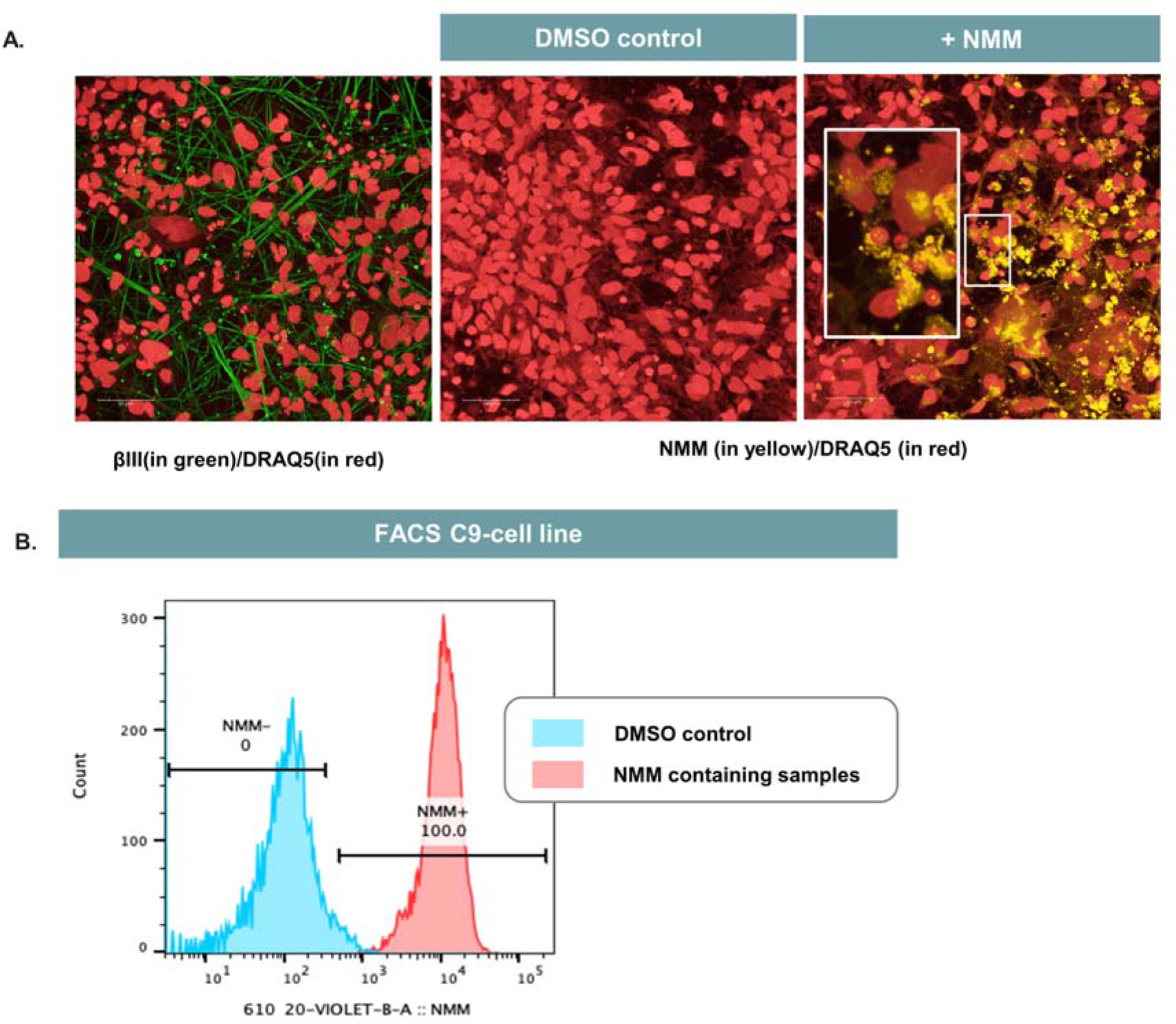
C9orf72 mutant iPSC patient derived spinal motor neurons show efficient fluorescent staining of RNA aggregates via the G4-binder NMM. **A. Confocal images of IPSC patient derived neurons.** The micrographs show the images taken by overlapping the Nuclear Stain DRAQ5 (in red) and the NMM channel (in yellow). In the DMSO control, no autofluorescence in cells can be observed in the NMM emission channel. Upon staining with the dye, the C9orf72 expansion containing neurons present fluorescent putative RNA aggregates. In the panel on the left, cells were stained with DRAQ5 and the neuronal marker β-tubulin III (in green). **B. Fluorescence-activated cell sorting (FACS) analysis for the C9-mutant motor neurons.** Cells treated with DMSO showed no characteristic fluorescence in the NMM-emission channel via FACS. NMM treatment resulted in detection of fluorescence of the characteristic NMM spectrum, indicating efficient incorporation into motor neurons.

## Discussion

Although unimolecular G4s are increasingly recognised drug targets for many diseases ranging from cancer and Cockayne Syndrome to Fragile X syndrome^28^, the biological relevance of mG4s has only recently started to be investigated^62^. Indeed, there is growing evidence of how mG4-formation could aid long-range interactions at enhancer-promoter regions in the genome^63^ and how mG4s could promote formation of phase-separated biomolecular condensates and/or aggregates ^10,62^. Furthermore, the recent discovery of an endogenous human protein that selectively binds mG4s over unimolecular structures strongly suggests that these multimeric species could be formed in cells in physiology and/or pathophysiology^32^.

Although unimolecular species have a kinetic advantage and therefore form more readily, mG4s are usually thermodynamically favoured in the crowded conditions of the cell^64^. However, despite the increasingly recognised biological role of mG4s, there is no specific tool to test for their presence *in vitro* or *in vivo*. Much like when unimolecular G4s were first discovered and there were no specific probes that would distinguish them from dsDNA in live cells, none of the currently used G4-specific probes (such as PDS^65^, PhenDC3^66^, BG4^67^ and NMM^48^) have been thoroughly studied in the context of mG4s, making their differentiation from unimolecular G4s in live-cell experiment challenging.

In this study we showed how (GGGGCC)*_n_* can arrange into mG4s *via* a combination of biophysical tools: circular dichroism demonstrating the formation of G4-structures, agarose gel electrophoresis confirming the formation of multimolecular species, and both NMM staining and KCl titrations providing further evidence to support the formation of these species in C9orf72 mutation-related repeats in ALS/FTD. We further observed a clear correlation between mG4-formation and the ability of (GGGGCC)_n_ to form biomolecular condensates *in vitro*. Since the pathological aggregate formation in ALS and FTD is strongly correlated with the presence of (GGGGCC)_n_ at high repeat numbers, we investigated the relationship between the observed nucleic acid-driven aggregation and repeat length itself. Importantly, we were able to show that higher repeat lengths require lower concentrations for aggregation to occur, which may bear significance for the disease setting. The repeat lengths used for this study are significantly below the recognised pathological threshold (*n* = 24)^8^, which is possibly why we only observe aggregation in the micromolar range. However, given our data, it is possible to infer that higher repeat lengths would aggregate at lower concentrations, potentially reaching biologically relevant ranges (a human cell contains ∼ 10–30 pg total RNA and 6 pg of DNA^68^).

The aggregates obtained were amorphous and solid in appearance. Although pathological condensation in ALS and FTD are often referred to as “liquid-like”^69,70,71^, it is reasonable to assume that additional protein components and RNAs^72^ in ALS/FTD-affected cells could modify the physical state of the aggregates. In fact, two of the main proteins implicated in ALS/FTD (TDP-43 and FUS) have shown significant G4-binding abilities^40,73^, suggesting that perhaps the protein could act in conjunction with G4-triggered aggregation in a live cell environment.

The condensates responded to staining with the G4-specific dye NMM and were successfully disassembled *via* guanine photo-oxidation when NMM was irradiated, which could be further explored as an approach to disassemble the pathological aggregates in live cells. As a proof of concept, the biological relevance of our study was assessed by testing the equivalent RNA (GGGGCC)*_6_* sequence, successfully confirming the G4-driven aggregation behaviour.

Furthermore, we demonstrated how the G4-ligand PDS can diminish mG4-induced aggregation of (GGGGCC)_10_ by driving the system to a lower molecularity state. In previously reported studies, G4-ligands have been shown to prevent aggregation events in ALS/FTD phenotypes^26,24^. It was hypothesised that the ligands interfere with the downstream biological functions of the gene, specifically in relationship to the toxic translated protein^52^, or displace proteins bound to the repeats. Although these pathways are still plausible, here we offer additional insights into the aggregation properties of such repeat expansion in the absence of proteins, demonstrating that PDS could also be affecting disease phenotypes in this context by shifting G4 towards lower molecularity species.

However, it is noteworthy that previous studies report conflicting relationships between the frequency of RNA foci and age of disease onset^74,75^ and no correlation between RNA foci aggregation and cognitive decline^76^ suggesting that the relationship between RNA foci and the clinical manifestation of C9orf72 ALS may be nuanced, context-specific, and perhaps less linked to cognitive function. Although our data show the potential relevance of mG4s in *C9orf72*, further work needs to be performed to tackle the underlying mechanism of action of pathological progression.

To further demonstrate the physiological relevance of our findings, we then incubated our mG4-based aggregates in the presence of the most widely recognized pathologically relevant protein in ALS/FTD (TDP-43). We revealed that both the protein and the nucleic acid aggregation properties are enhanced when incubated together. Furthermore, we confirmed the physiological relevance of our observation, by successfully staining C9orf72 mutation carrying patient-derived iPSC-motor neurons with NMM. NMM staining has revealed the presence of G4-structures in pathological aggregates, suggesting that the model described in this study could be of both pathophysiological and therapeutic relevance.

In summary, our data indicate a protein-free pathway for the formation of RNA condensates that could be relevant for the initiation and/or progression of C9orf72 mutation-related ALS and FTD pathogenesis, making mG4s a potential therapeutic target for these devastating and currently incurable neurodegenerative diseases. Importantly, this does not exclude the relevance of proteins or other canonical and non-canonical nucleic acid structures as contributors to these aggregates, as indicated by the condensation synergy observed in the presence of TDP-43.

Currently, the only three FDA-approved drugs that have shown a modest increase in survival rate in randomised clinical trials across the majority of ALS cases are Riluzole, Edaravone and AMX0035 ^77,78,79,80^ but the need for clinically impactful therapies is apparent and urgent. Other trials targeting specific familial causes of ALS include a promising GAPmer antisense oligonucleotide (ASO) for SOD1 mutation carrying patients (approximately 2% of all ALS cases) recently being approved by the FDA^81^. However, the first C9orf72 ASO (BIIB078) trial was discontinued by Biogen for failure to meet any of its secondary endpoints (NCT03626012). Although further trials using different and possibly more effective C9orf72 ASOs are underway, it is crucial to consider emerging insights into the disease mechanisms to maximise our repertoire for therapeutic strategy. So far, candidate drugs have been developed to target protein–led aggregation but have shown only partial effectiveness^82^, suggesting the need to explore alternative approaches. A recent study investigating a small molecule with the ability to cross the blood-brain barrier and bind to r(GGGGCC)*_n_* promoting formation of a hairpin structure over G4s, which could offer the starting point for a new therapeutic pathway^25^. Altogether, our findings in the context of these aforementioned studies strongly suggest that targeting RNA structures within the (GGGGCC)*_n_* repeat represents an exciting and potentially tractable strategy for treating ALS and FTD and is worthy of further investigation.

## Methods

### 1. (GGGGCC)_n_ mG4s annealing protocol

(GGGGCC)*_n_* (*n* = 1-12) was purchased from IDT in its lyophilised form and diluted in MilliQ water to reach a stock concentration of 1 mM. The stocks were heated to 40 ℃ for 5 minutes to ensure complete solvation. The DNA samples were then further diluted to the desired annealing concentration (500-0 µM) in 10 mM TRIS-HCl buffer at pH 7.4 (filtered with 0.2 µm sterile syringe filters) and 30% poly(ethylene glycol) BioUltra 200 (PEG) was added to the clear solution. The samples were vortexed and subjected to a first denaturing step in a C1000 Touch™ Thermal Cycler (heating to 95 ℃ for 1h followed by cooling to 25 ℃ with a temperature ramp of 0.25 ℃/min). 500 mM KCl was added to the denatured solution and a further heating cycle was applied (heating to 95 ℃ for 1h followed by cooling to 25 ℃ with a temperature ramp of 0.02 ℃/min). To note is that after the long annealing procedure, significant evaporation is observed in the samples. The phase diagram was developed by reducing evaporation as much as possible to estimate an accurate concentration. For this purpose, the author recommends using an annealing volume of > 30 µL. For other applications where exact concentration is not of interest smaller volumes can also be adopted.

Annealing in absence of PEG was performed in the same way but with a slower cooling rate (heating to 95 ℃ for 1h followed by cooling to 25 ℃ with a temperature ramp of 0.01 ℃/min).

### 2. Circular Dichroism spectroscopy protocol

Pre-annealed (GGGGCC)*_n_* (n =1-12) DNA samples were diluted to 5 µM in 10 mM TRIS-HCl buffer pH 7.4 with a final volume of 150 µL. The samples were transferred in Quartz High Precision cell (1 mm light path) and analysed in the Jasco J-715 Spectropolarimeter. The parameters used for each run were the following: Sensitivity (100 mdeg); Wavelength start and end (320 nm – 220 nm); Data Pitch 0.5 nm with continuous scanning mode and scanning speed of 100 nm/min; Response (2 sec); Band Width (2.0 nm); Accumulation (3).

### 3. Agarose gel electrophoresis protocol SYBR safe staining

Pre-annealed (GGGGCC)*_n_* (n =1-12) DNA samples were diluted to 10 µM in 10 mM TRIS-HCl buffer pH 7.4 with a final volume of 12 µL. 2 µL of Purple Dye were added to the solution to visualise the samples. The agarose gel matrix was prepared by mixing 1.5 g of agarose in 50 mL 1X TBE buffer and 3 µL SYBR safe stain, heated by microwaving and left to polymerise for 20 minutes prior to running. The samples were run at 65 V for 85 mins at room temperature or 50 V for 150 minutes on ice (depending on the application) and imaged with Typhoon FLA 9500 on the EtBr channel. Running the gel on ice for longer times and at a lower voltage makes it easier to distinguish multimolecular species.

#### NMM staining

Pre-annealed (GGGGCC)*_n_* (n =1-12) DNA samples were diluted to 10 µM in 10 mM TRIS-HCl buffer pH 7.4 with a final volume of 12 µL. 2 µL of 50% glycerol were added to the solution to visualise the samples. The agarose gel matrix was prepared by mixing 1.5 g of agarose in 50 µL 1X TBE buffer, heated by microwaving and left to polymerise for 20 minutes prior to running. The samples were run at 65 V for 85 mins at room temperature or 50 V for 150 minutes on ice (depending on the application). The gel was then incubated in a solution of 50 µL 2mM NMM and 50 mL 1X TBE for 1h covered to avoid photobleaching of the dye. The gel was washed once with 1X TBE for 10 minutes and imaged with Typhoon FLA 9500 on the EtBr channel. Running the gel on ice for longer times and at a lower voltage makes it easier to distinguish multimolecular species. Notably, due to the overlapping emission/excitation range of NMM and SYBR safe, both stainings cannot be performed on the same gel.

### 4. Imaging via confocal microscopy Phase diagram

Leica SP8 Inverted Confocal microscope (HC PL APO CS2 10x/0.40 DRY | HC PL APO CS2 20x/0.75 DRY objectives) was used for the imaging. In order to define aggregation on the phase diagram, pre-annealed (GGGGCC)*_n_* (n =1-12) DNA samples were directly transferred in PDMS wells and imaged by irradiating the sample with the 458 nm excitation laser on the brightfield channel.

#### FAM-labelled samples

Leica SP8 Inverted Confocal microscope (HC PL APO CS2 10x/0.40 DRY | HC PL APO CS2 20x/0.75 DRY objectives) was used for the imaging. Pre-annealed samples labelled with 2-50% FAM were diluted to 10 µM in 10 mM TRIS-HCl buffer pH 7.4, imaged by irradiating with the 488 nm laser and with an emission filter from 510-550 nm.

#### NMM-stained samples

Leica SP8 Inverted Confocal microscope (HC PL APO CS2 10x/0.40 DRY | HC PL APO CS2 20x/0.75 DRY objectives) was used for the imaging. Pre-annealed samples were diluted to 10 µM in 10 mM TRIS-HCL buffer pH 7.4 and irradiated with the 405 nm laser. 10 µM NMM was then added to the sample and left to incubate for 10 minutes in a dark environment. The sample was then imaged on the emission filter 600-650 nm.

#### G4-ligands containing samples

Leica SP8 Inverted Confocal microscope (HC PL APO CS2 10x/0.40 DRY | HC PL APO CS2 20x/0.75 DRY objectives) was used for the imaging. Pre-annealed (GGGGCC)_10_ in presence of the 10 µM PDS was directly transferred in PDMS wells and imaged by irradiating the sample with the 458 nm excitation laser on the brightfield channel. As PDS is solubilised in presence of 0.1% DMSO, a DMSO control was also added to the experiment.

#### RRM1–2 containing samples

Leica SP8 Inverted Confocal microscope (HC PL APO CS2 10x/0.40 DRY | HC PL APO CS2 20x/0.75 DRY objectives) was used for the imaging. Pre-annealed (GGGGCC)_11_ in presence of the 10 µM RRM1–2 was directly transferred in PDMS wells and imaged by irradiating the sample with the 405 nm excitation laser on the brightfield channel. NMM was then added in a 1:1 NMM:DNA ratio and left to incubate for 10 minutes. The sample was then imaged again on the emission filter 600-650 nm.

#### (GGGGCC)_11_-FAM and RRM1–2-ALEXA633 samples

Leica SP8 Inverted Confocal microscope (HC PL APO CS2 10x/0.40 DRY | HC PL APO CS2 20x/0.75 DRY objectives) was used for the imaging. Pre-annealed (GGGGCC)_11_ (10% fluorescently labelled with FAM) in presence of the 10 µM RRM1–2 (fluorescently labelled with ALEXA633) was directly transferred in PDMS wells and firstly imaged by irradiating the sample with the 405 nm excitation laser on the brightfield channel. The DNA component was imaged by irradiating with the 488 nm laser and with an emission filter from 510-550 nm. The protein component was imaged by irradiating with the 633 nm laser and with an emission filter 610-700 nm).

#### NMM-photooxidation

Leica SP8 Inverted Confocal microscope (HC PL APO CS2 10x/0.40 DRY | HC PL APO CS2 20x/0.75 DRY objectives) was used for the imaging. Pre-annealed (GGGGCC)*_6_* sample was diluted to approximately 10 µM in 10 mM TRIS-HCL buffer pH 7.4 and irradiated with the 405 nm laser. 10 µM NMM was then added to the sample and left to incubate for 10 minutes in a dark environment. A portion of the sample was then irradiated at 405 nm at 100% for 1 minute and complete disassembly of that portion of the aggregate was observed both in brightfield or on the emission filter 600-650 nm.

#### FRAP (Fluorescence recovery after photobleaching)

Leica SP8 Inverted Confocal microscope (HC PL APO CS2 10x/0.40 DRY | HC PL APO CS2 20x/0.75 DRY objectives) was used for the imaging. Pre-annealed samples labelled with 10% FAM were imaged by irradiating with the 488 nm laser and with an emission filter from 510-550 nm. A region of the well was irradiated at 100% power intensity for 30 seconds until no fluorescence was observed. Recovery after photobleaching was monitored every five minutes for 30 minutes.

### 5. dsDNA condensates annealing protocol

Condensates made from dsDNA nanostars were prepared by mixing four nanostar core strands (N4_core1, N4_core2, N4_core3, N4_core4) and two linker strands (L_AA1 and L_AA2) in a TrisHCl buffer containing 500mM KCl at a final nanostar concentration of 1 μM and linkers concentration of 2 μM. A volume of 70μL was then loaded into borosilicate glass capillaries (inner dimensions: 0.4 mm × 4 mm × 50 mm, CM Scientific) with a micropipette. Prior to loading, capillaries were cleaned by sonication in 1% Hellmanex III (HellmaAnalytics) at 60°C for 15 minutes. The surfactant was removed through five rounds of rinsing with deionised (DI) water, followed by a round of sonication in ultrapure water (Milli-Q). After loading the sample, the capillary ends were capped with mineral oil and sealed with epoxy glue (Araldite) on a coverslip. The sample was annealed by incubating at 95℃ for 30 minutes and cooled down between 85℃ and 25℃ (-0.1℃/min cooling rate). Samples were transferred from the glass capillary to a PDMS well for imaging. All strands were purchased from Integrated DNA Technologies (IDT) and reconstituted in TE. Sequences are provided in the SI, **Figure S4** (A).

### 6. (GGGGCC)_n_ mG4s annealing protocol in presence of PDS

(GGGGCC)*_10_* was purchased from IDT in its lyophilised form and diluted in MilliQ water to reach a stock concentration of 1 mM. The stock was heated to 40 ℃ for 5 minutes to ensure complete solvation. The DNA sample was then further diluted to the desired annealing concentration (150 µM) in 10 mM TRIS-HCl buffer at pH 7.4 (filtered with 0.2 µm sterile syringe filters) and 30% poly(ethylene glycol) BioUltra 200 (PEG) was added to the clear solution. The samples were vortexed and subjected to a first denaturing step in a C1000 Touch™ Thermal Cycler (heating to 95 ℃ for 1h followed by cooling to 25 ℃ with a temperature ramp of 0.25 ℃/min). 500 mM KCl and 10 µM PDS were added to the denatured solution and a further heating cycle was applied (heating to 95 ℃ for 1h followed by cooling to 25 ℃ with a temperature ramp of 0.02 ℃/min). To solubilise the PDS, 0.1% DMSO was also added to the solution, therefore a DMSO control has also been added to the results. To note is that after the long annealing procedure, significant evaporation is observed in the samples. The phase diagram was developed by reducing evaporation as much as possible to estimate an accurate concentration. For this purpose, the author recommends using an annealing volume of > 30 µL.

### 7. (GGGGCC)_11_ and RRM1–2 aggregates RRM1–2 expression and purification

Purification of the TDP-43 RRMs fragment was carried out as previously described^57^. Briefly, the protein construct was expressed in a pET-SUMO expression vector containing the kanamycin antibiotic resistance gene. The plasmid was expressed by heat-shock transformation in BL21 *Escherichia coli* (*E. Coli*) as a protein fused with a SUMO solubilization tag and a 6×His tag. Cells were grown in Luria-Bertani (LB) with 50 μg/mL kanamycin at 37 °C with 200 rpm shaking until they reached an OD of 0.7 at 600 nm. Protein expression was induced with 0.5 mM IPTG, and cells were grown overnight at 18 °C with 200 rpm shaking. Cells were then collected by centrifugation at 4000 rcf for 20 min at 4°C and resuspended in lysis buffer (10 mM potassium phosphate buffer pH 7.2, 150 mM KCl, 5mM imidazole, 5% v/v glycerol, 1mg/ml lysozyme, complete^TM^ EDTA-free Protease Inhibitor tablet by Roche, 1 μg/ml DNase I and 1 μg/ml RNaseA). The resuspended cells were sonicated on ice for 20 minutes (15s on and 45s off pulses, 20% amplitude) and soluble proteins were recovered by centrifugation at 18,000 rcf for 45 min at 4°C. The supernatant was filtered using a 0.22 μm filter and loaded onto a HisTrap HP 5 ml column (Cytiva) which was pre-equilibrated with 10 mM potassium phosphate buffer pH 7.2 supplemented with 15 mM imidazole (binding buffer). The 6xHis-SUMO-construct was eluted with phosphate buffer supplemented with 300 mM imidazole (elution buffer). After measuring the concentration of RRMs on a Nanodrop (Thermo Fisher Scientific), the eluate was dialysed overnight at 4 °C against phosphate buffer in the presence of Tobacco Etch Virus (TEV) protease (1:10 TEV:protein construct molar ratio) to remove the 6xHis-SUMO tag. A second nickel-affinity chromatography was performed, and the flowthrough was loaded onto a HiTrap Heparin column (Cytvia) which was pre-equilibrated with 10 mM phosphate buffer. The protein was eluted with high-salt phosphate buffer (10 mM potassium phosphate buffer pH 7.2, 1.5 M KCl) and applied on a HiLoad 16/60 Superdex 75 prep grade column (Cytvia) pre- equilibrated with 10 mM phosphate buffer for size-exclusion chromatography. Protein concentration was taken on a Nanodrop, and protein identity and purity were checked by PAGE and mass spectrometry. All protein purification steps were carried out using an AKTA Pure (Cytvia).

#### SDS-PAGE

Samples were prepared in 4X LDS sample buffer and 10X reducing agent and boiled at 95°C for 5 min. Samples were run on 4-12% Bis-Tris NuPAGE gels (Thermo Fisher Scientific) at 180 V for 35 min. Gels were stained with InstantBlue Coomassie Protein Stain (Abcam) for 15 mins, washed in water for 30 mins and imaged on an ImageQuant LAS 4000.

#### Electrospray ionization mass spectrometry (ESI-MS)

ESI-MS was performed on purified protein samples (50 μM) buffer-exchanged into HPLC grade water to confirm molecular weight and sample purity. ESI-MS was performed by Malgorzata Puchnarewicz using the Chemistry Mass Spectrometry facilities available at the Molecular Sciences Research Hub, Department of Chemistry, Imperial College London.

#### RRM1–2 labelling with ALEXA 633

Purified RRMs were fluorescently tagged using Alexa Fluor™ 633 C5 Maleimide (Thermo Fisher Scientific). The dye was added to a sample of purified protein (1:10 protein:dye molar ratio) in 10 mM potassium phosphate buffer pH 7.2 and incubated at 25 °C for 4 hours on a roller mixer with slow rotation. The solution was then dialysed against 10 mM potassium phosphate buffer pH 7.2 overnight to remove unbound dye.

#### DNA/protein aggregates formation

(GGGGCC)_11_ was annealed at different concentrations (300, 100 and 1 µM as per mG4- annealing protocol. The samples were then incubated for 3 days at 37C with 10 µM of TDP- 43 RBD.

#### Transmission electron microscopy (TEM)

To prepare TEM grids, 10 μL of sample was pipetted on Formvar/Carbon coated 300 mesh copper grids (Agar Scientific) and left to incubate for 2Cmins. Excess sample was removed with Whatman filter paper. The grids were then washed with dH2O and stained withCuranyl acetateC(2% w/v) for 2 mins. Excess uranyl acetateCwas removed, and the grid was left to dry for at least 5 mins. For each sample, images were taken in at least 5 locations on the grid to ensure consistent structures were present across the grid. Grids were measured on a T12 Spirit electron microscope.

### 8. iPSC derived neuronal cell culture

Experimental protocols were carried out according to approved regulations and guidelines by UCL Hospitals National Hospital for Neurology and Neurosurgery and UCL Institute of Neurology joint research ethics committee (09/0272). iPSCs were maintained with Essential 8 Medium media (Life Technologies) on Geltrex (Life Technologies) at 37°C and 5% carbon dioxide. iPSCs were passaged when reaching 70% confluency using EDTA (Life Technologies, 0.5 mM). iPSC cultures underwent differentiation into spinal cord motor neurons as previously described^59^. Briefly, iPSCs were plated to 100% confluency and then differentiated to a spinal neural precursor fate by sequential treatment with small molecules, day 0–7: 1 μM Dorsomorphin (Tocris Bioscience), 2 μM SB431542 (Tocris Bioscience), and 3.3 μM CHIR99021 (Tocris Bioscience), day 7–14: 0.5 μM retinoic acid (Sigma Aldrich) and 1 μM Purmorphamine (Sigma Aldrich), day 14–18: 0.1 μM Purmorphamine. After neural conversion and patterning, 0.1 μM Compound E (Bio-techne) was added for terminal differentiation into spinal cord motor neurons.

#### Immunocytochemistry (ICC)

Cells were seeded on clear bottom 96 well plates (Falcon) at a density of 30,000 cells per well. After 24 hours, Compound E (Bio-techne) was added for terminal differentiation into spinal cord motor neurons. At day 6 of terminal differentiation, DRAQ5 (Merck) was applied to the cells diluted at 1:1000 in media and incubated for 10 minutes to stain nuclei. Cells were then fixed in 4% paraformaldehyde for 15-20 minutes at room temperature, followed by permeabilization using 0.3% Triton-X (NMM and antibody) and blocking with 5% BSA in PBS for 60 minutes (antibody). Either 10 µM NMM (Cambridge Bioscience) or 0.5% DMSO diluted in 0.15% Triton-X containing PBS, or a primary antibody was then applied. The NMM/DMSO was incubated for 40 hours at 4°C while the primary antibody anti-tubulin beta-III (#802001, BioLegend), diluted at 1:2000 in 0.15% Triton-X in PBS, was left overnight at 4°C. Plates treated with NMM/DMSO were imaged directly. Secondary antibody incubation was then performed using Alexa Fluor-conjugated secondary antibody 568 nm (anti-rabbit) at 1:1000 dilution in PBS at room temperature. Cells were imaged with Opera Phenix (PerkinElmer) at 40X, Z series of images were used. Standard imaging and acquisition settings were applied. All cell images were processed to better visualization with a 20% increase in brightness and contrast.

#### Flow cytometry

Motor neurons were dissociated into single cells using Accutase (Life Technologies). The cell suspensions were fixed using 4% paraformaldehyde for 15-20 minutes and then permeabilised using 0.3% Triton-X in PBS for 1 hour at room temperature. Subsequently, they were treated with either 10 µM NMM (Cambridge Bioscience) or 0.5% DMSO diluted in 0.15% Triton-X overnight at 4°C. The samples (10,000 cells/sample) were examined using a BD LSRFortessa analyser operated by the FACSDiva software (BD), and the results were analysed using the FlowJo software.

## Acknowledgements

Graphics were designed using Procreate design app and ChemDraw. The authors thank the Crick Flow Cytometry and High Throughput Screening STPs for their technical assistance.

## Authors contributions

FR developed the experimental protocols and conducted all experiments unless otherwise stated. DT designed and produced the dsDNA condensates control. AH expressed and purified TDP-43 and contributed to performing the TDP-43 incubation experiments. JL cultured the iPSC patient derived neuronal cells, aided with imaging for the NMM staining experiment and performed FACS. TM synthesised PDS and supported the related experiments. LM supported the RNA and DNA photooxidation experiments. RRS helped generating the 3D reconstructed image of the fluorescence condensates. FAA and DMV aided in the design of the TDP-43 experiments. DMV and AH performed the TEM experiments. RB, RP, MPH, YW and MDA contributed to designing the cellular experiments. FR analysed all the data with support from MDA, LDM and RP. FR and MDA co-wrote the paper, with support from LDM and RP. MDA and FR has designed the overall research design with substantial support by LDM and YE. MDA, LDM and YE supervised the research. All authors discussed the results and edited the paper.

## Funding

This work was supported by the following funding bodies:

MDA is supported by Biotechnology and Biological Sciences Research Council (BBSRC) David Phillips Fellowship [BB/R011605/1]; MDA is a Lister Institute Fellow and is supported by a Lister Institute Research Prize. FR is supported by a Leverhulme Trust, Leverhulme Cellular Bionics scholarship [EP/S023518/1]; LDM is supported by Royal Society University Research Fellowship [UF160152, URF\R\221009] and Royal Society Research Fellows Enhanced Research Expenses [RF/ERE/210029], also supporting RRS. LDM, RRS, DT and LM are supported by the European Research Council (ERC) under the Horizon 2020 Research and Innovation Programme [ERC-STG No 851667 NANOCELL]. YE is supported by United Kingdom Research and Innovation Future Leaders Fellowship [MR/S031537/1]. TM is supported by [EPSRC G98210]. FAA is supported by United Kingdom Research and Innovation Future Leaders Fellowship [MR/S033947/1] and Alzheimer’s Research UK [ARUK-PG2019B-020]. RP holds an MRC Senior Clinical Fellowship (MR/S006591/1) and a Lister Research Prize Fellowship. AH is supported by The Engineering and Physical Research Council (Grant Number: EP/S023518/1). RB is an NIHR Academic Clinical Lecturer in Neurology at UCL and is funded by an Academy of Medical Sciences Starter Grant for Clinical Lecturers (SGL027\1022). MPH is funded through an MRC grant (MR/S006591/1). YW is supported by a Crick i2i grants (P2022-0007).

Facility for Imaging by Light Microscopy (FILM) at Imperial College London is part-supported by funding from the Wellcome Trust [grant 104931/Z/14/Z] and BBSRC [grant BB/L015129/1].

Funding for open access charge: Imperial College London.

## Materials and Correspondence

For further information refer to corresponding authors Dr Di Antonio, Dr Di Michele and Prof. Rickie Patani. Tel: +44 (0)20 7594 5866; (MDA) | (LDM) and (RP)

## Notes

### Competing Interest Statement

The authors have declared no competing interest.

### Summary of Updates

This is a revised version of the manuscript following the first round of revision after submission.

