## Supplementary Information for "The ALS/FTD-related C9orf72 hexanucleotide repeat expansion forms RNA condensates through multimolecular G-quadruplexes"

**Figure S1: A. (GGGGCC)$_n$ FAM labelled AGE** – FAM labelled (GGGGCC)$_n$ was annealed at 1 µM under G4-forming conditions (KCl 500 mM). Three distinct bands can be detected, respectively corresponding to bi-molecular and tetra-molecular species, with higher molecular weight species formed at higher repeat lengths, while lower molecularity species are more prominent at lower repeat lengths. **B. (GGGGCC)$_n$ FAM labelled AGE** – FAM labelled (GGGGCC)$_n$ was annealed at 350 µM under G4-forming conditions (KCl 500 mM). No distinct bands can be observed due to sample aggregation but is evident how higher repeat lengths lead to higher molecularity species.

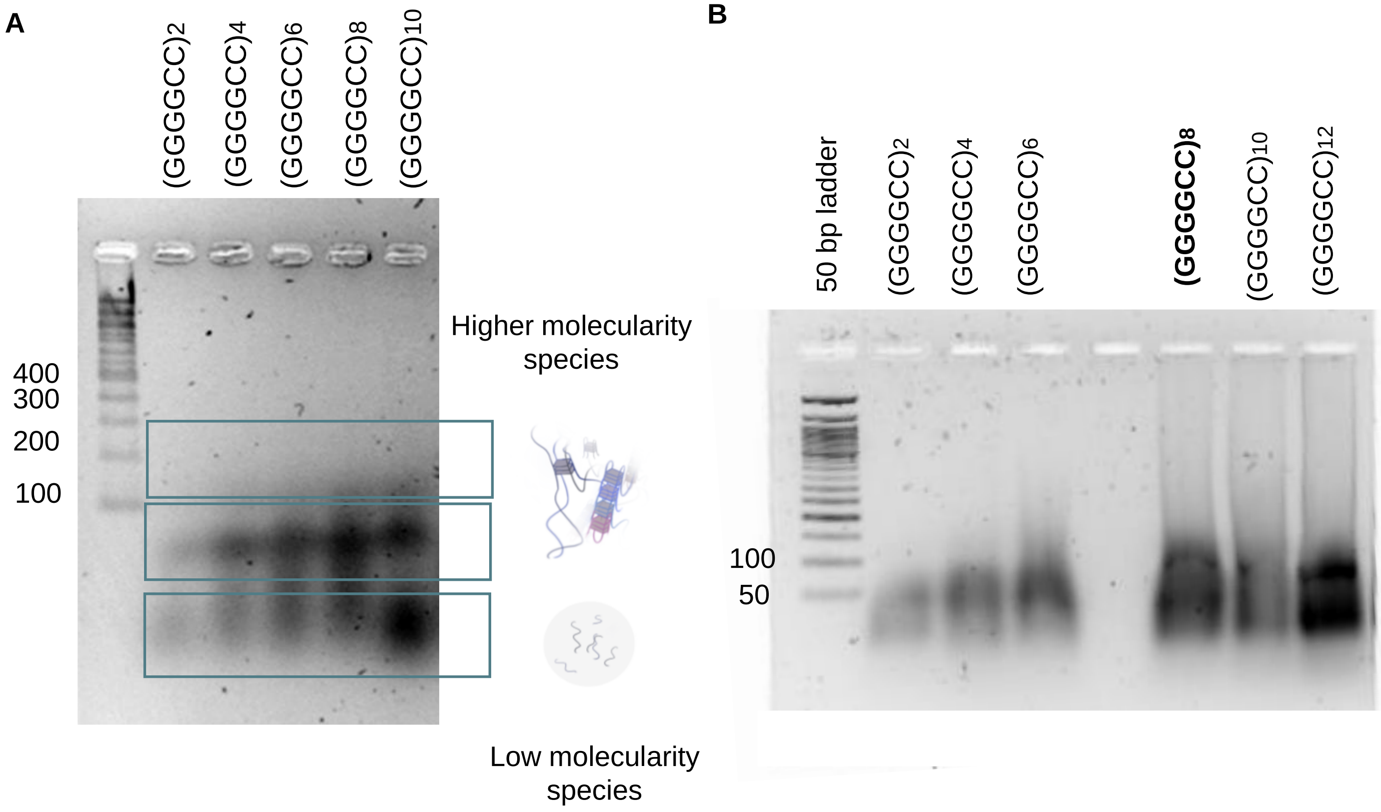

**Supplementary Information**

**Figure S2: A. (GGGGCC)$_{6}$ FAM staining experiment** – (GGGGCC)6 was annealed (50% FAM labelled) in mG4s forming conditions. Brightfield and fluorescence channel overlayed. **B. (GGGGCC)6 FAM staining experiment fluorescence images**– (GGGGCC)6 was annealed (50% FAM labelled) in mG4-forming conditions. **C. (GGGGCC)$_{10}$ FRAP experiment.** – (GGGGCC)10 was annealed (10% FAM labelled) in mG4-forming conditions. Upon laser excitation FAM labelled strands photobleached, leading to a local reduction of fluorescence intensity. The photobleached strands did not diffuse within the reset of the aggregate after 30 minutes, indicating a solid-like state. 10 µm scalebar.

**A.**

**GGGGCC*6* – FAM labelled**

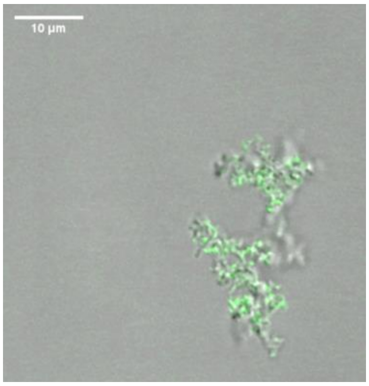


**B.**

**GGGGCC*6* – FAM labelled**

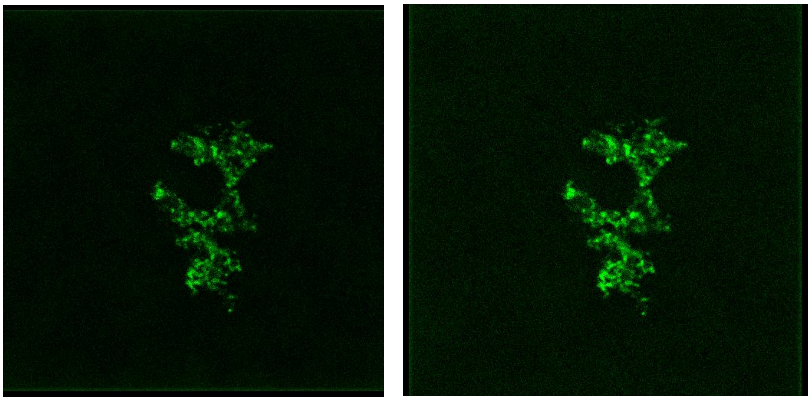

**C.**

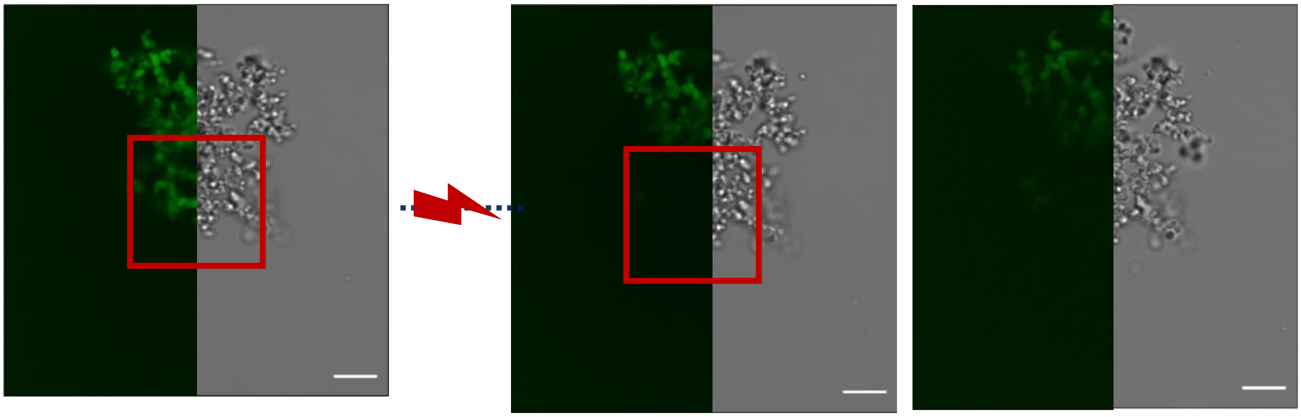

Before irradiation

After irradiation

30 mins after irradiation

**Figure S3:** Phase Diagram with brighfield microscopy images.

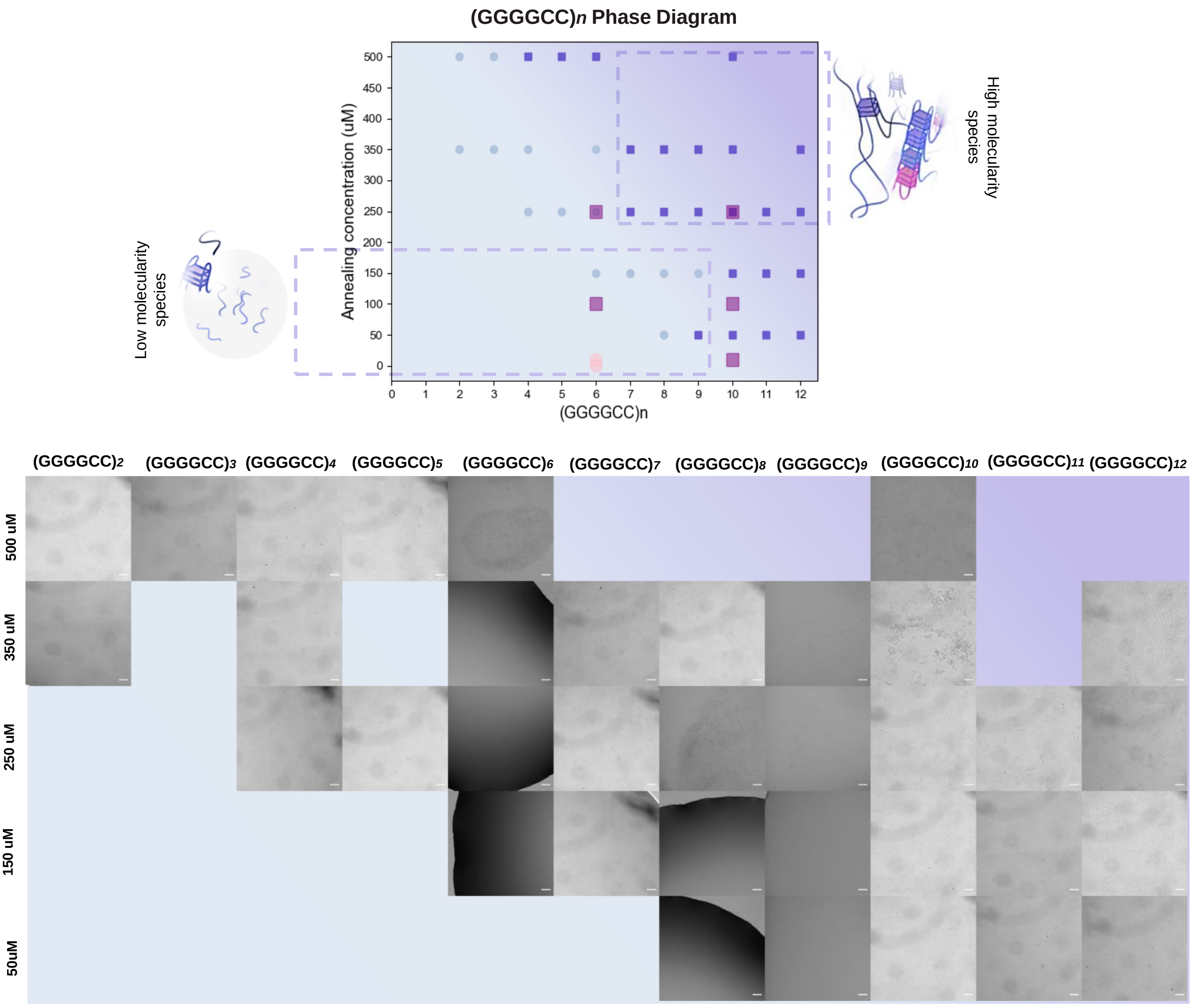

**Figure S4: dsDNA condensates (nanostars) control for NMM staining and photooxidation experiments - A.** Sequences used to build DNA nanostars. **B.** Brightfield imaging and NMM fluorescent channel of the dsDNA condensates. No NMM fluorescence is observed in presence of dsDNA. **C.** dsDNA nanostar control for the photooxidation experiment. 10 µm scalebar.

**A.**

| Name | Sequence (5’-3’) |
|---|---|
| N4_ core1 | **GATCGC**CGCCGCAATCACGCGCGTGCTCGGCGCCAGCAGTCCTGGCG |
| N4_core2 | **GATCGC**CGCCAGGACTGCTGGCGCCGTCGCTTCTCTTCATAACAACG |
| N4_core3 | **GATCGC**CGTTGTTATGAAGAGAAGCGTCGCTCTGGCACAGGTGTACG |
| N4_core4 | **GATCGC**CGTACACCTGTGCCAGAGCGTGCACGCGCGTGATTGCGGCG |
| L_AA1 | **GCGATC**CGCAAACCAGCAAGCTCACG |
| L_AA2 | **GCGATC**CGTGAGCTTGCTGGTTTGCG |

**B.**

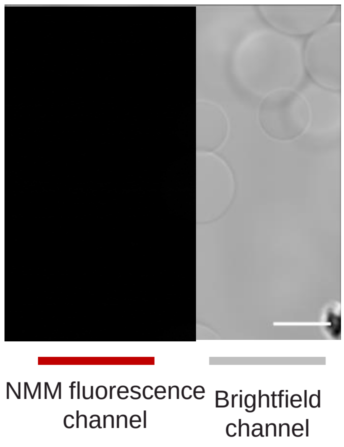


**C.**

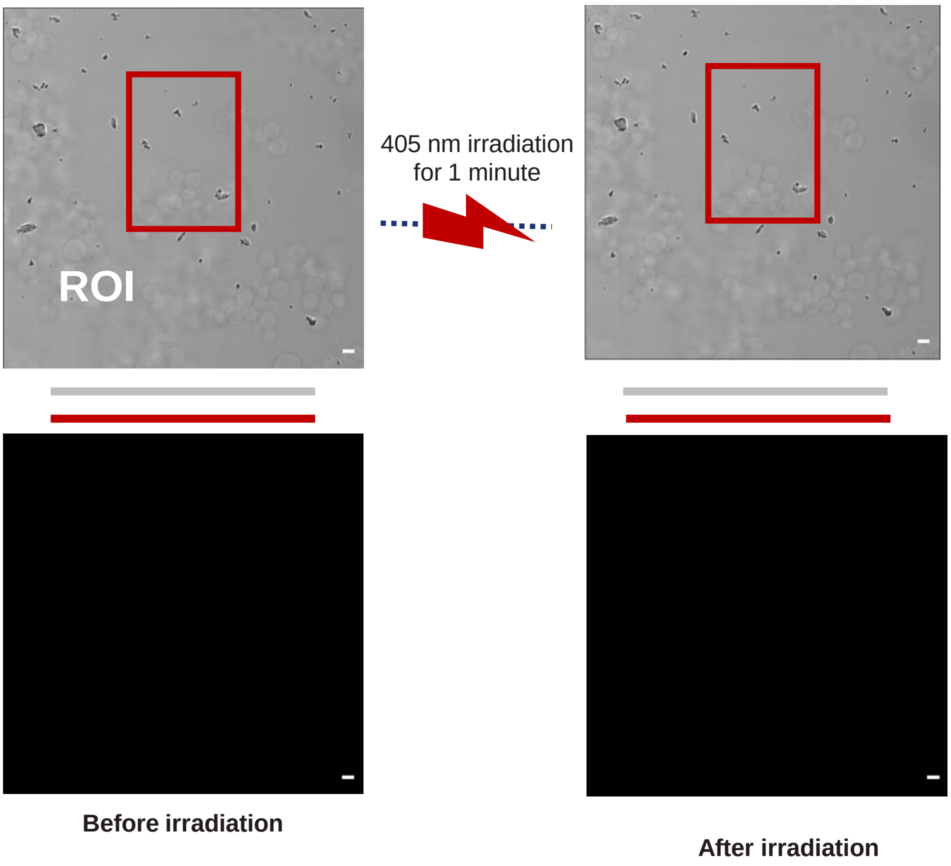

**Figure S5: DNA (GGGGCC)$_{11}$ aggregates form at physiological concentrations of KCl.** Brightfield imaging in of 500 µM (GGGGCC)$_{11}$ annealed with 100/300/500 mM KCl. 50 µm scalebar.

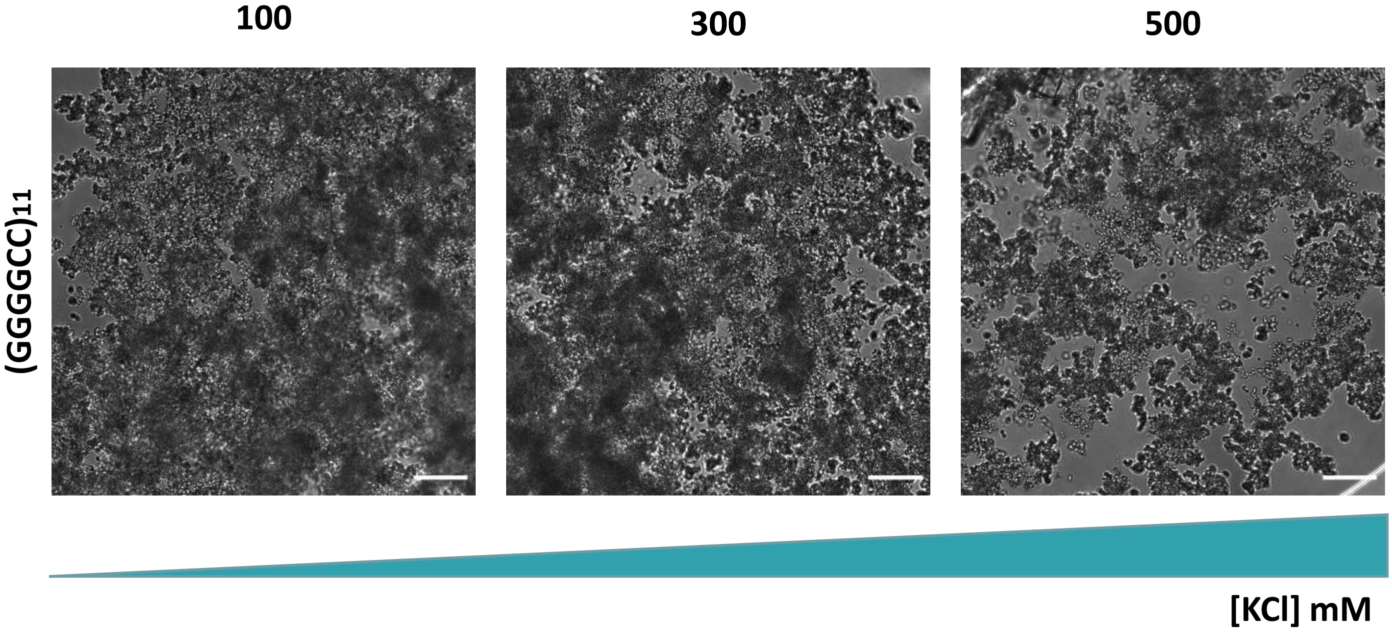


**Figure S6: DNA (GGGGCC)$_{11}$ aggregates form in absence of the crowding agent PEG.** Brightfield imaging of (GGGGCC)$_{11}$ annealed under mG4-forming conditions at 500 µM. On the right, a faster cooling rate is used in presence of the PEG crowding agent. On the left, a slower cooling rate in absence of PEG. (See methods for details) 50 µm scalebar.

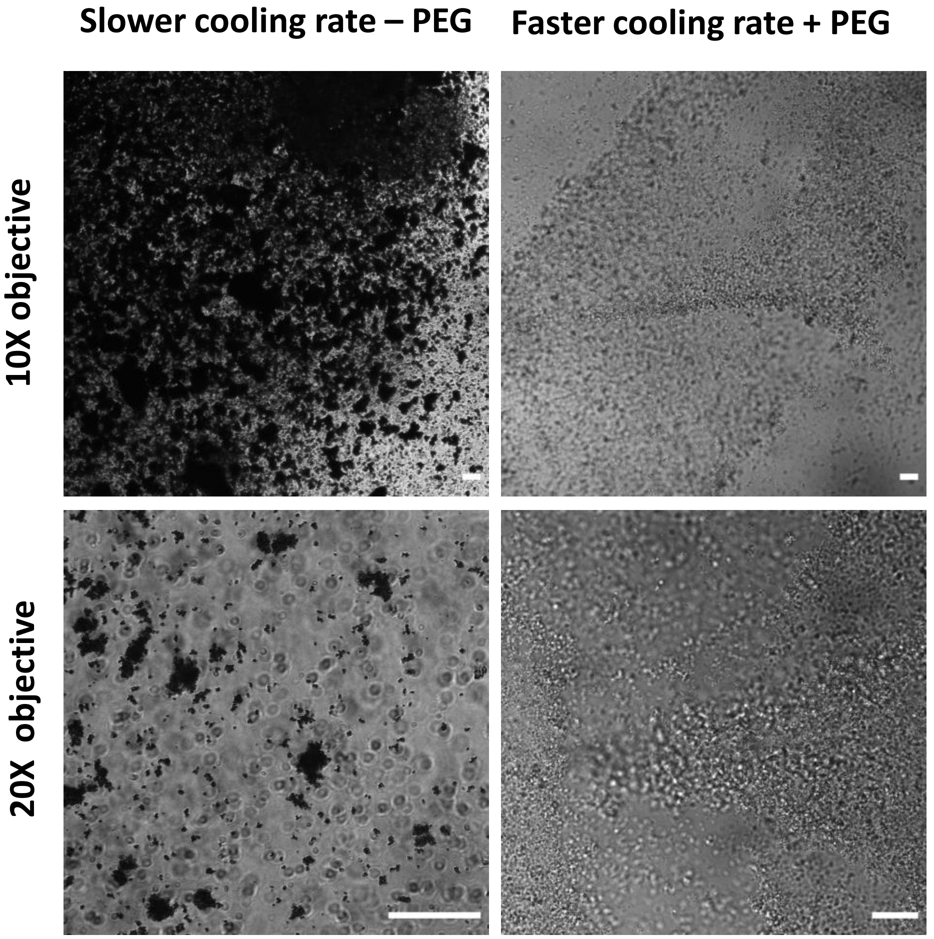

**Figure S7: A. G4-ligand experiment agarose gel electrophoresis – SYBR safe stain**. $(GGGGCC)_{10}$ was annealed under mG4-forming conditions. Control samples in absence of PDS present higher molecularity species that appear as smearing in the gel. **B. G4-ligand experiment agarose gel electrophoresis – NMM stain**. $(GGGGCC)_{10}$ was annealed under mG4-forming conditions. All samples containing G4-forming DNA appear stained by the probe.

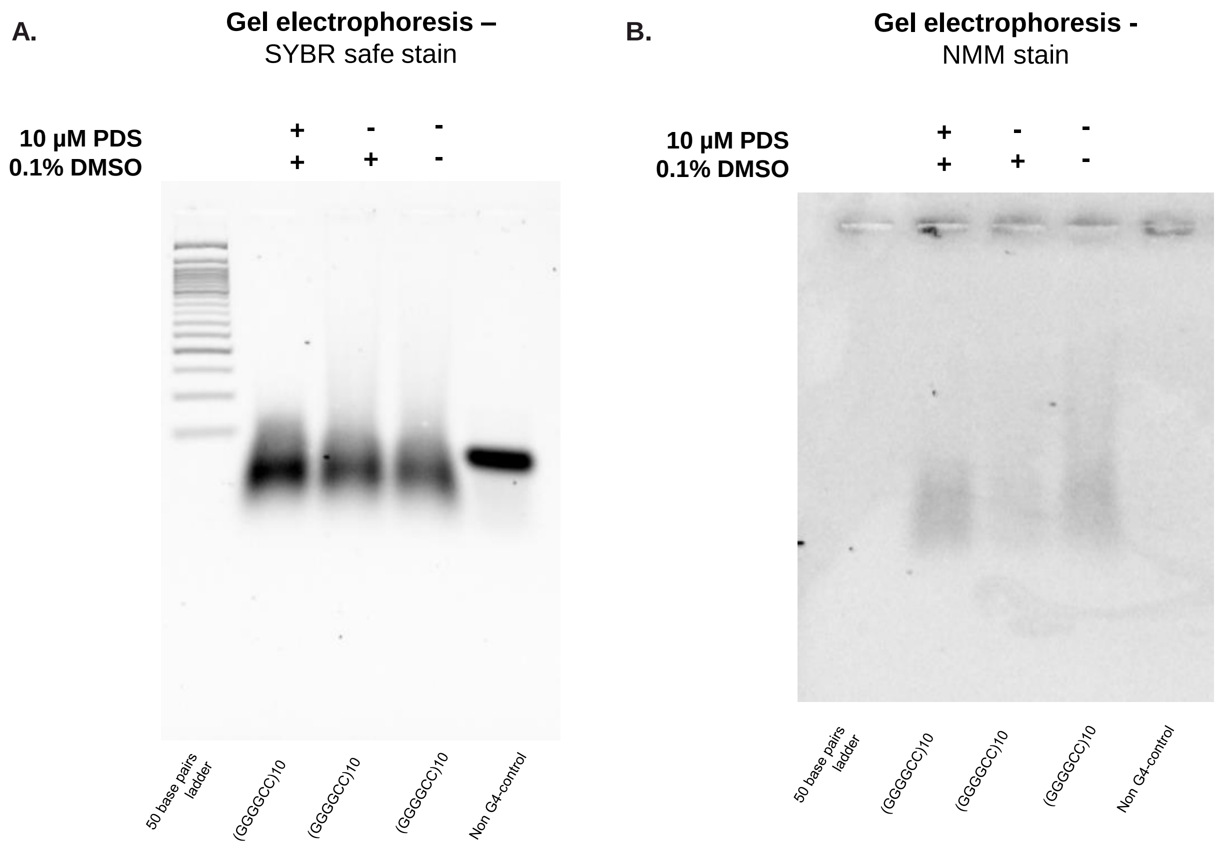


**Figure S8: A. r(GGGGCC)n aggregates.** Brightfield images of $r(GGGGCC)_n$ annealed at different concentrations.

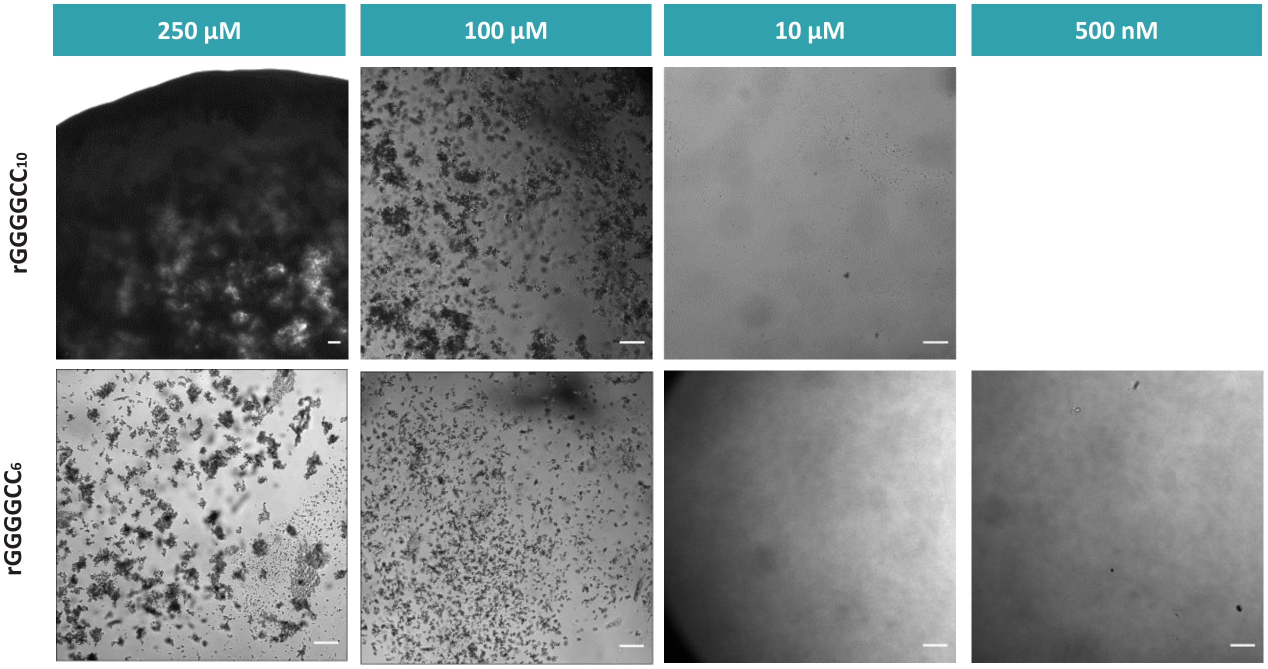

**Figure S9: TDP-43 RRM1-2 after purification. A.** SDS-PAGE gel and expected molecular weight form amino acid sequence. **B**. ESI-MS spectra after deconvolution.

**A.**

### SDS-PAGE of TDP-43 RRM1-2 post size-exclusion chromatography (SEC)

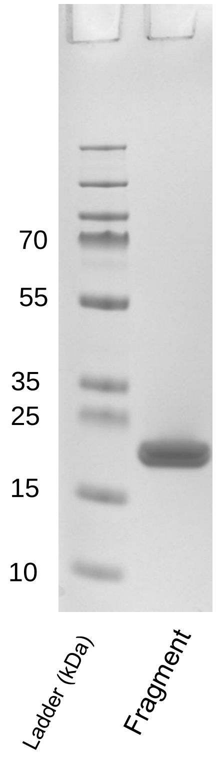


**Amino acid sequence of RBDs:**

KTSDLIVLGLPWKTTEQDLKEYFSTFGEVLMVQVKKDLKTGHSKG
FGFVRFTEYETQVKVMSQRHMIDGRWCDCKLPNSKQSQDEPLR
SRKVFVGRCTEDMTEDELREFFSQYGDVMDVFIPKPFRAFAFVTF
ADDQIAQSLCGEDLIIKGISVHISNAEPKHNSNRQ

Average M.W.: 19396.89219 ±1.31502

Monoisotopic M.W.: 19384.64622

**B.**

### ESI-MS of TDP-43 RRM1-2 post size-exclusion chromatography (SEC)

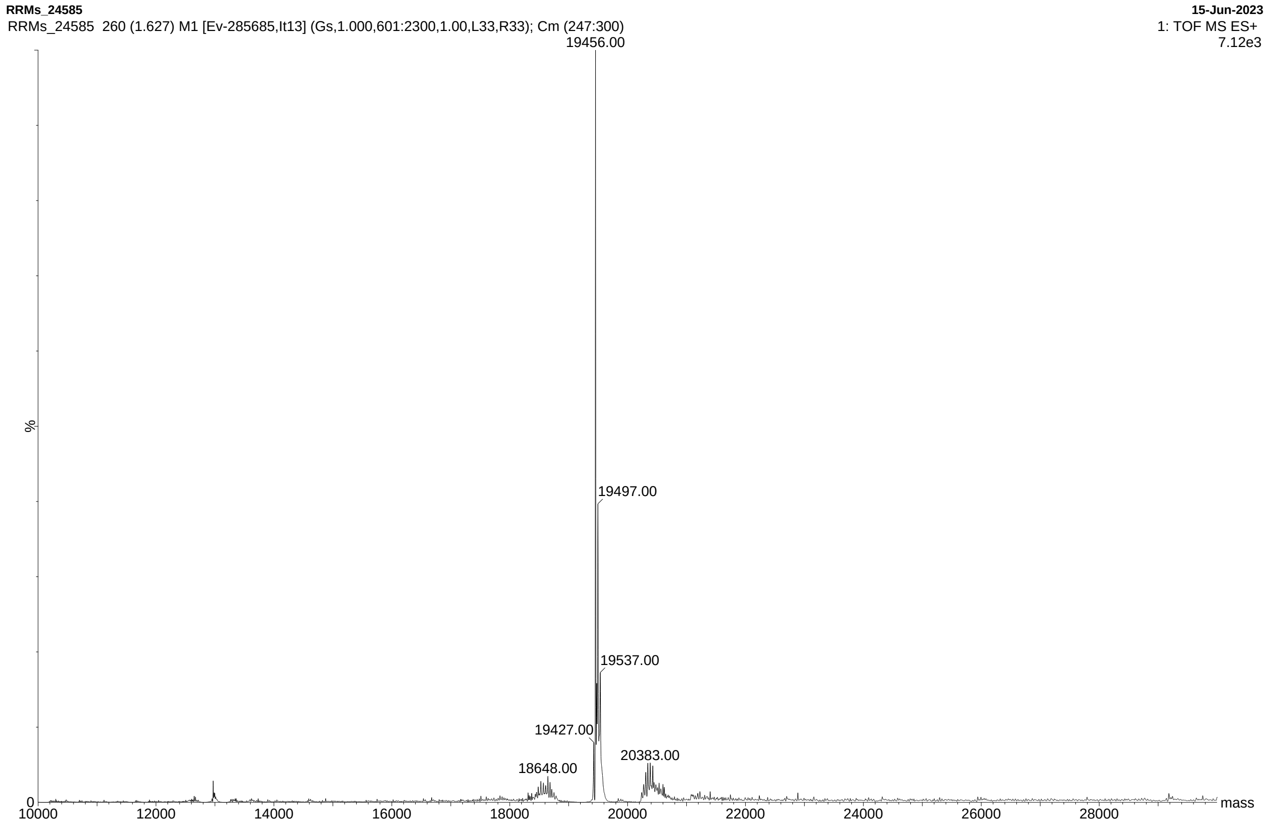

**Figure S10: A. (GGGGCC)11 and TDP-43 aggregates.** Confocal images of (GGGGCC)11 annealed at 100 μM (FAM label) and incubated for three days with 10 μM TDP-43 RBD (Alexa 633 label). B. **(GGGGCC)11 and TDP-43 aggregates NMM-stained.**

**A.**

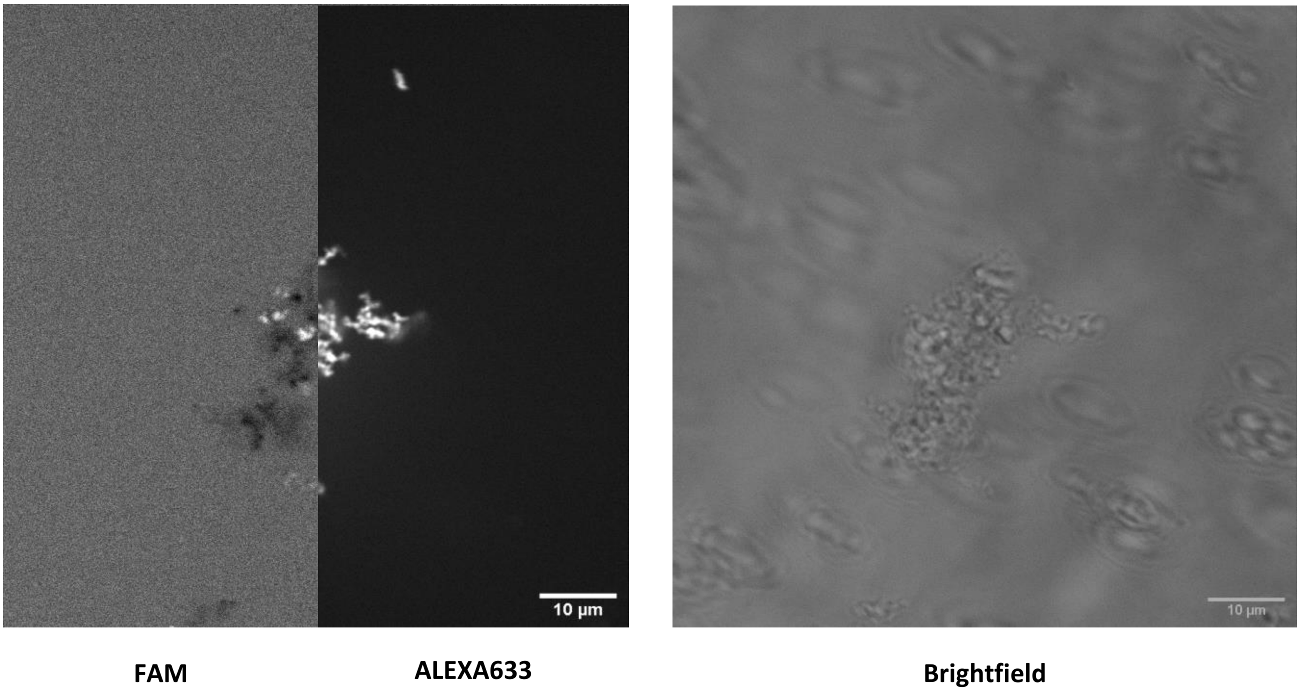


**B.**

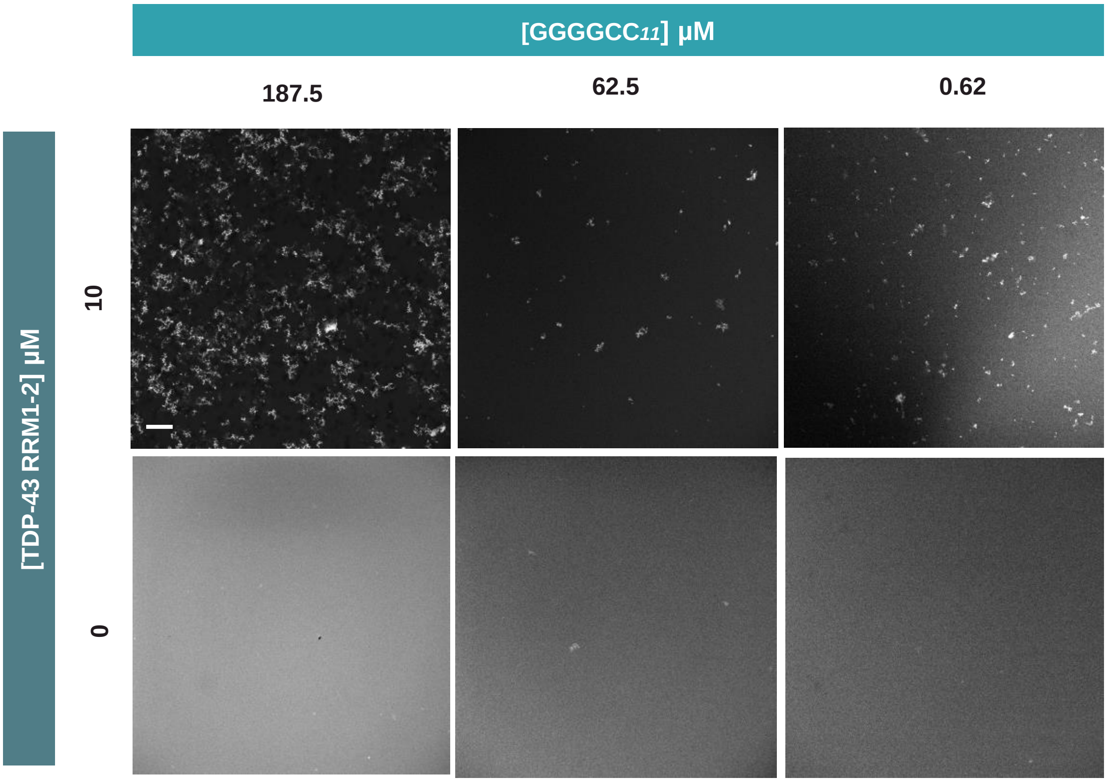

**Figure S11: TEM images TDP-43 RRM1-2 aggregates.** TEM images confirm that smaller aggregates can be seen beyond the resolution of a confocal microscope. These aggregates are still larger in presence of the DNA component.

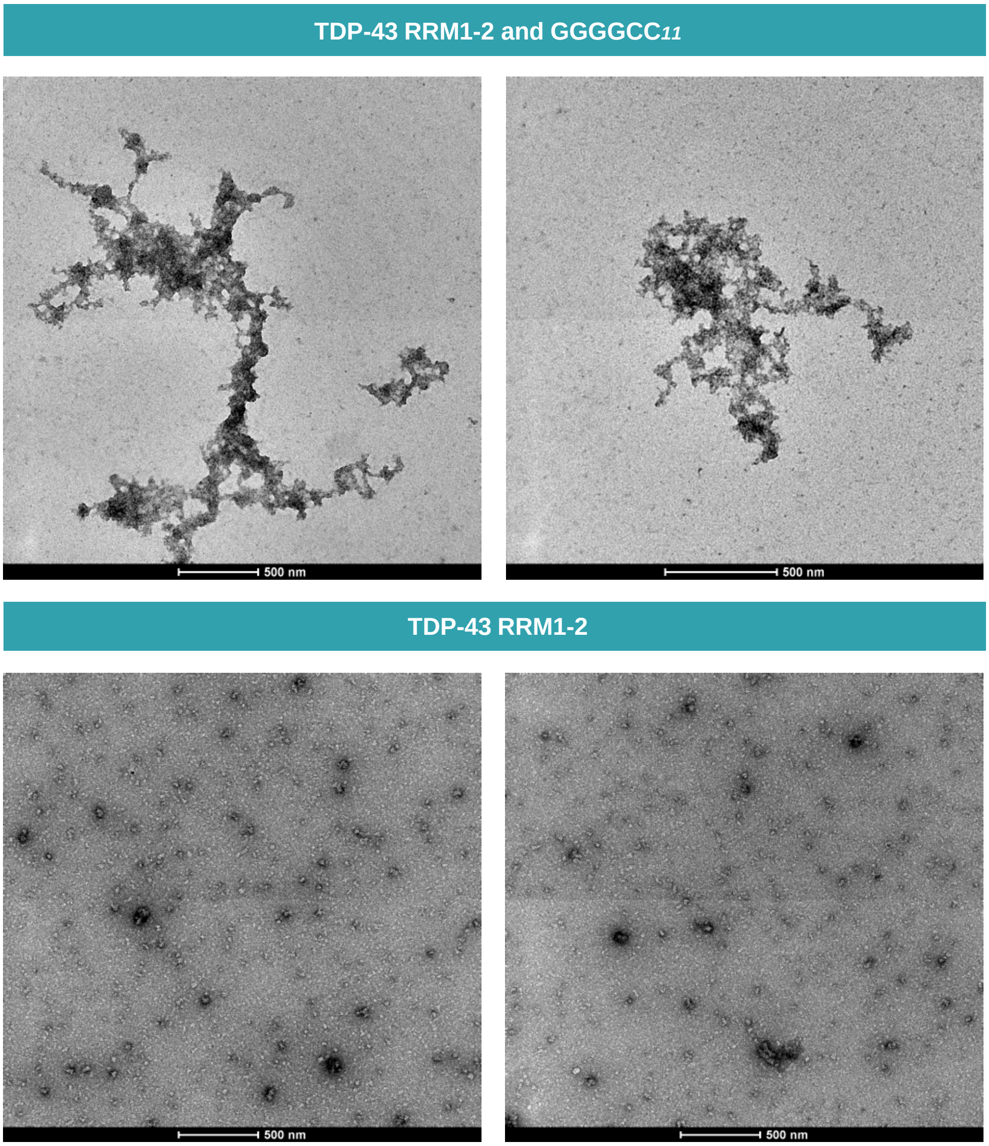

**Figure S12: FACS data for cell-lines stained with NMM.** Cells fluorescing in the NMM-fluorescence channel were counted in the C9-cell line and healthy neuronal cells controls. The DMSO controls show that most cells are not fluorescing the NMM channel – indicating no significant autofluorescence effect.

### FACS data

#### C9-cell line

**DMSO control**

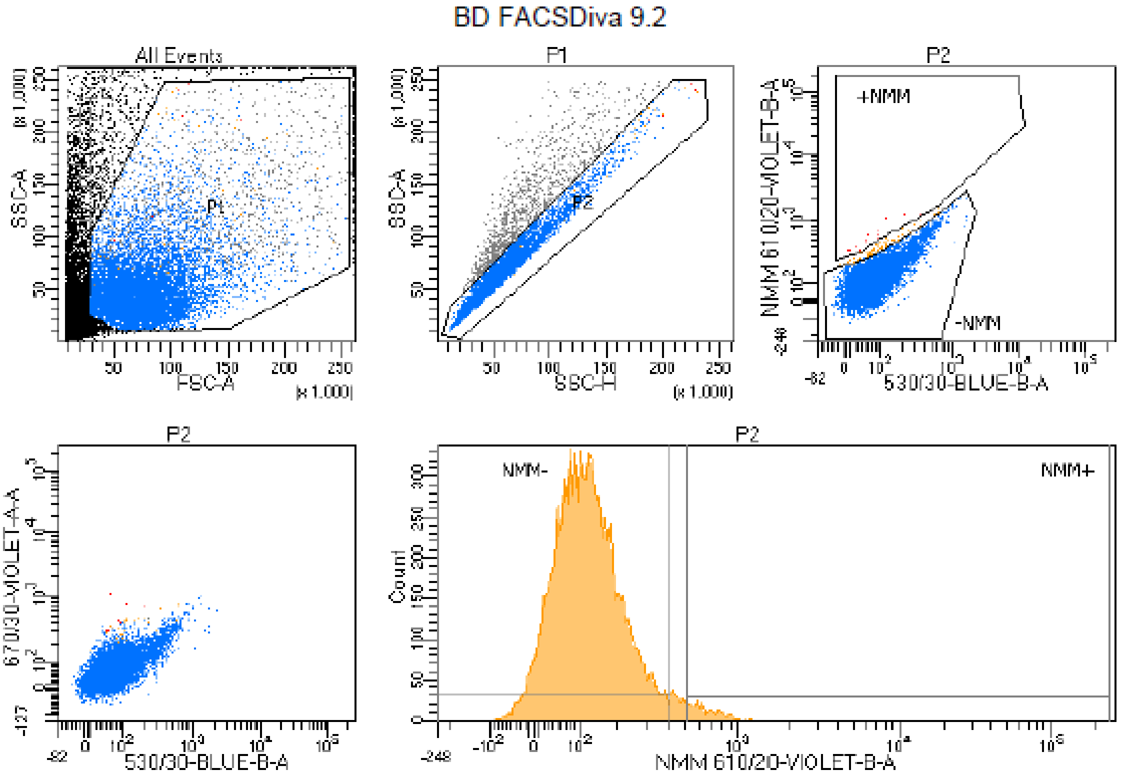


Tube: N6 DMSO

| Population | #Events | %Parent | %Total |
|---|---|---|---|
| All Events | 21,263 | #### | 100.0 |
| P1 | 11,154 | 52.5 | 52.5 |
| P2 | 10,000 | 89.7 | 47.0 |
| NMM+ | 292 | 2.9 | 1.4 |
| NMM- | 9,512 | 95.1 | 44.7 |
| +NMM | 13 | 0.1 | 0.1 |
| -NMM | 9,919 | 99.2 | 46.6 |

| Population | NMM 610/20-VIOLET-B-A Median | 670/30-VIOLET-A-A Median |
|---|---|---|
| P2 | 111 | 60 |
| NMM+ | 657 | 301 |
| NMM- | 106 | 57 |
| +NMM | 532 | 305 |
| -NMM | 110 | 60 |

**NMM – containing sample**

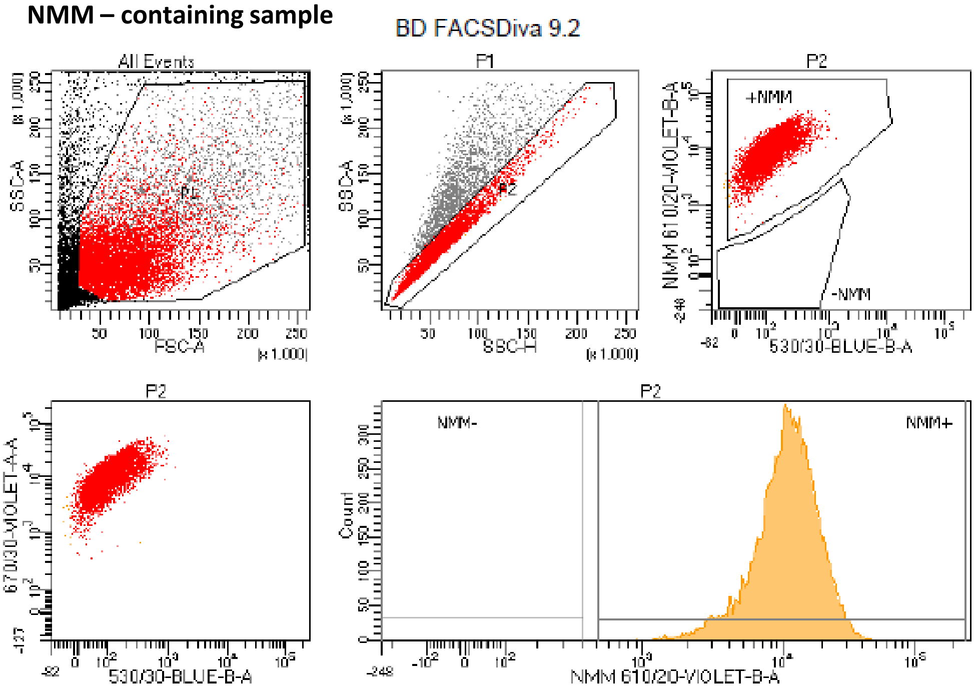


Tube: N6 10uM NMM

| Population | #Events | %Parent | %Total |
|---|---|---|---|
| All Events | 19,631 | #### | 100.0 |
| P1 | 12,152 | 61.9 | 61.9 |
| P2 | 10,000 | 82.3 | 50.9 |
| NMM+ | 9,998 | 100.0 | 50.9 |
| NMM- | 1 | 0.0 | 0.0 |
| +NMM | 9,987 | 99.9 | 50.9 |
| -NMM | 0 | 0.0 | 0.0 |

| Population | NMM 610/20-VIOLET-B-A Median | 670/30-VIOLET-A-A Median |
|---|---|---|
| P2 | 10,414 | 11,870 |
| NMM+ | 10,415 | 11,871 |
| NMM- | 373 | 429 |
| +NMM | 10,423 | 11,879 |
| -NMM | #### | #### |
